# Mutations in *C. elegans* neuroligin-like *glit-1*, the apoptosis pathway and the calcium chaperone *crt-1* increase dopaminergic neurodegeneration after 6-OHDA treatment

**DOI:** 10.1101/203067

**Authors:** Sarah-Lena Offenburger, Elisabeth Jongsma, Anton Gartner

**Affiliations:** School of Life Sciences, University of Dundee, Dundee DD1 5EH, UK

## Abstract

The loss of dopaminergic neurons is a hallmark of Parkinson’s disease, the aetiology of which is associated with increased levels of oxidative stress. We used *C. elegans* to screen for genes that protect dopaminergic neurons against oxidative stress and isolated *glit-1 (<u>gli</u>o<u>t</u>actin (Drosophila neuroligin-like) homologue)*. Loss of the *C. elegans* neuroligin-like *glit-1* causes increased dopaminergic neurodegeneration after treatment with 6-hydroxydopamine (6-OHDA), an oxidative- stress inducing drug that is specifically taken up into dopaminergic neurons. Furthermore, *glit-1* mutants exhibit increased sensitivity to oxidative stress induced by H_2_O_2_ and paraquat. We provide evidence that GLIT-1 acts in the same genetic pathway as the previously identified <u>t</u>etra<u>sp</u>anin TSP-17. After exposure to 6-OHDA and paraquat, *glit-1* and *tsp-17* mutants show almost identical, non-additive hypersensitivity phenotypes and exhibit highly increased induction of oxidative stress reporters. TSP-17 and GLIT-1 are both expressed in dopaminergic neurons. In addition, the neuroligin-like GLIT-1 is expressed in pharynx, intestine and several unidentified cells in the head. GLIT-1 is homologous, but not orthologous to neuroligins, transmembrane proteins required for the function of synapses. The *Drosophila* GLIT-1 homologue Gliotactin in contrast is required for epithelial junction formation. We report that GLIT-1 likely acts in multiple tissues to protect against 6-OHDA, and that the epithelial barrier of *C. elegans glit-1* mutants does not appear to be compromised. We further describe that hyperactivation of the SKN-1 oxidative stress response pathway alleviates 6-OHDA-induced neurodegeneration. In addition, we find that mutations in the canonical apoptosis pathway and the calcium chaperone *crt-1* cause increased 6-OHDA-induced dopaminergic neuron loss. In summary, we report that the neuroligin-like GLIT-1, the canonical apoptosis pathway and the calreticulin CRT-1 are required to prevent 6-OHDA-induced dopaminergic neurodegeneration.

**Author summary:** Neurons use dopamine as a chemical messenger to mediate diverse behaviours. The gradual loss of dopaminergic neurons in specific brain areas is a hallmark of Parkinson’s disease. The increased occurrence of highly reactive oxygen radicals, also called oxidative stress, is assumed to contribute to the demise of dopaminergic neurons. In this study we searched for genes that protect dopaminergic neurons against oxidative stress. We used the nematode *C. elegans*, a well-characterised model organism whose dopamine signalling system is very similar to that of humans. When *C. elegans* is exposed to 6-hydroxydopamine, an oxidative stress-inducing compound, dopaminergic neurons gradually die. Our major findings include: (i) absence of the neuroligin-like gene *glit-1* causes highly increased cell death of dopaminergic neurons after 6-OHDA exposure; (ii) GLIT-1 acts in a similar manner as the previously identified tetraspanin TSP-17; (iii) GLIT-1 and TSP-17 also protect *C. elegans* from other types of oxidative stress; and (iv) the programmed cell death pathway and a calcium chaperone protect dopaminergic neurons as well. Collectively, this study shows that apoptosis proteins, the calcium chaperone CRT-1 and the neuroligin-like GLIT-1 protect against neurodegeneration after oxidative stress exposure.

## Introduction

Dopamine acts as a neurotransmitter to control attention, cognition, motivation and movement [1]. The progressive loss of dopaminergic neurons in the substantia nigra of the midbrain is a hallmark of Parkinson’s disease, the second most common neurodegenerative disorder (for review [2,3]). The etiology of the disease is largely unknown and there is no causative treatment for Parkinson’s disease to date (for review [4]). Less than 10% of Parkinson’s patients report a positive family history while the majority of instances are sporadic [3]. Most Parkinson’s disease cases appear to be evoked by the interplay of one or multiple genetic defects and environmental factors [4]. A major risk factor for both genetic and sporadic forms of Parkinson’s disease is oxidative stress, the imbalance between the production of reactive oxygen species and the antioxidant response (for review [5]). Reactive oxygen species can damage cellular components and ultimately cause cell loss [5]. However, it is not clear which pathways mediate dopaminergic cell death after oxidative stress exposure (for review [6]).

The gradual loss of dopaminergic neurons can be modelled by exposure to the oxidative stress- inducing drug 6-hydroxydopamine (6-OHDA) (for review [7]). 6-OHDA is an hydroxylated dopamine analogue which is specifically taken up into dopaminergic neurons by the dopamine transporter and which inhibits complex I of the mitochondrial electron transport chain [7]. The resulting formation of reactive oxygen species is assumed to trigger 6-OHDA-induced neurodegeneration [7]. 6-OHDA was found in human brain samples [8] and in higher concentrations in the urine of Parkinson’s patients [9].

6-OHDA exposure of *C. elegans* leads to the selective loss of dopaminergic neurons [10]. The components of dopaminergic synaptic transmission are highly conserved in *C. elegans* (for review [11]). The nematodes’ dopaminergic neurons are not required for viability and overt movement but they mediate subtle motor behaviours [11]. *C. elegans* hermaphrodites possess eight dopaminergic neurons, all of which have mechanosensory roles [12]: four CEP (<u>cep</u>halic sensilla) and two ADE (<u>a</u>nterior <u>de</u>irids) neurons in the head, and two PDE (<u>p</u>osterior <u>de</u>irids) neurons in the midbody. These neurons can be visualised by expression of a fluorescent protein from the promoter of the dopamine transporter [11]. Upon 6-OHDA exposure, dopaminergic neurons display blebbed processes, dark and rounded cell bodies and chromatin condensation, and eventually degenerate [10]. The pathways mediating 6-OHDA-induced dopaminergic cell death are unknown. Interference with the core apoptosis machinery does not prevent 6-OHDA-induced dopaminergic neurodegeneration; however, affected cells do not show typical morphological features for necrotic cell death either [10].

We used the *C. elegans* 6-OHDA model of dopaminergic neurodegeneration to screen for genes that protect from oxidative stress. Previously, this approach led to the characterisation of the tetraspanin TSP-17 which acts in dopaminergic neurons and alleviates 6-OHDA-induced neurodegeneration [13]. In this study, we report that the previously uncharacterised neuroligin-like gene *glit-1* prevents 6- OHDA-induced dopaminergic cell death. Furthermore, *glit-1* likely acts in the same genetic pathway as *tsp-17: glit-1* and *tsp-17* mutants are short-lived, exhibit similarly increased expression of oxidative stress reporters and show basically identical sensitivity upon exposure to oxidative stress-inducing compounds. We further found that the apoptosis pathway and the calcium chaperone CRT- 1 protect dopaminergic neurons from 6-OHDA-induced cell death.

## Results

### *gt1981* mutants exhibit highly increased dopaminergic neurodegeneration after 6-OHDA exposure

Animals carrying mutations in neuroprotective genes are expected to exhibit increased cell death when exposed to a low dose of a neurodegenerative drug. To find genes that prevent dopaminergic neuron death, a *C. elegans* strain with GFP-labelled dopaminergic neurons (Fig 1A) was mutagenised with ethyl methanesulfonate. The F2 progeny of these mutagenised animals was exposed to the neurodegenerative drug 6-hydroxydopamine (6-OHDA) at a dose that does not induce cell death in wild-type animals. We treated *C. elegans* synchronised at the first larval stage (L1), the first developmental phase after egg hatching at which the ADE and CEP dopaminergic head neurons are already fully differentiated. 72 hours after 6-OHDA treatment, the now adult population was screened for mutant animals exhibiting excessive loss of dopaminergic head neurons. We isolated the *gt1981* mutant line, which exhibited highly increased 6-OHDA-induced dopaminergic neurodegeneration when L1, L2 or L3 stage larvae were exposed to 6-OHDA (Fig 1B). Later stages were not severely affected by *gt1981* mutation (Fig 1B). We further found that the general healthiness of *gt1981* mutants is not compromised following 6-OHDA exposure as determined by measuring animal development: *gt1981* mutants exhibited slightly slower growth than wild-type animals, but this delay was not exacerbated following 6-OHDA treatment (S1A Fig). Thus, the *gt1981* mutation affects dopaminergic neuroprotection, but not the overall development of *C. elegans* larvae after 6-OHDA exposure.

**Fig 1.**
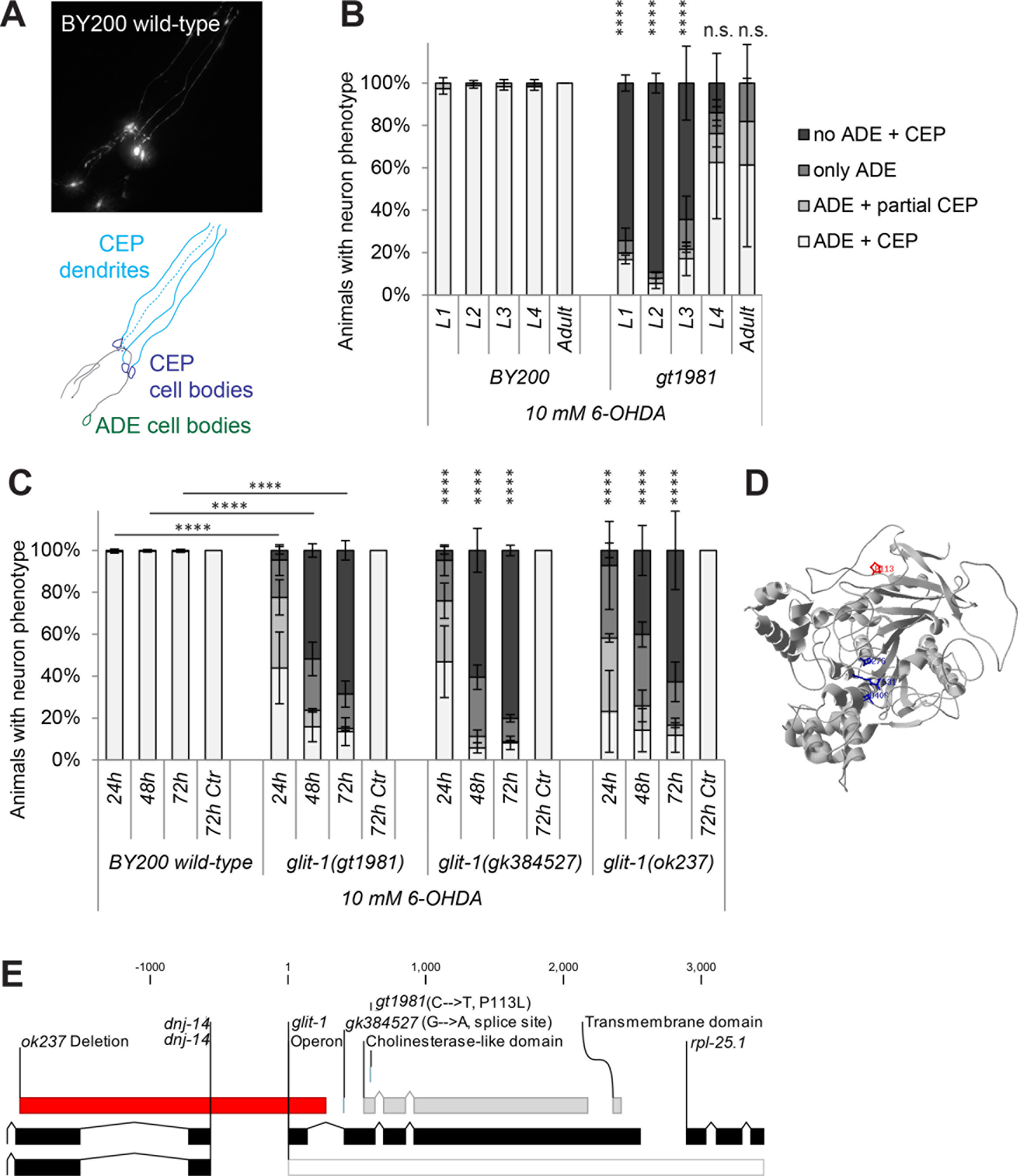
*glit-1(gt1981)* exhibits increased 6-OHDA-induced dopaminergic neurodegeneration. (A) GFP-labelled *C. elegans* dopaminergic head neurons – 4 CEP neurons and 2 ADE neurons – in BY200 wild-type animals. (B) Remaining dopaminergic head neurons in BY200 wild-type animals and *gt1981* mutants 48 hours after treating L1-L4 larval stages or adult animals with 10 mM 6-OHDA. Animals possessing all neurons were scored as ‘ADE + CEP’ (white bar), those with partial loss of CEP but intact ADE neurons as ‘ADE + partial CEP’ (light grey bar), those with complete loss of CEP but intact ADE neurons as ‘only ADE’ (dark grey bar) and those with complete loss of dopaminergic head neurons as ‘no ADE + CEP’ (black bar). Error bars = SEM of 2 biological replicates, each with 25-40 animals per stage and strain. Total number of animals per condition n = 50-80 (****p<0.0001, **p<0.01, n.s. p>0.05; G-Test comparing BY200 wild-type and mutant data of the same lifecycle stages). (C) Dopaminergic head neurons 24, 48 and 72 hours after treatment with 10 mM 6-OHDA and 72 hours after treatment with ascorbic acid only (‘72h Ctr’) for BY200 wild-type or *glit-1* mutant animals. Error bars = S.E.M. of 2 biological replicates for *glit-1(ok237)* and 3 biological replicates for all the other strains, each with 60-115 animals per strain and concentration. Total number of animals for the ‘72h Ctr’ experiment n = 30-100 and for all other conditions n = 130-340 (****p<0.0001; G- Test comparing BY200 wild-type and mutant data of the same time point). (D) GLIT-1 protein structure prediction based on homology modelling with acetylcholinesterase (PDB ID: 2W6C). The *gt1981* point mutation leads to a proline to glycine conversion (P113G) and is indicated in red. The amino acids replacing the acetylcholinesterase catalytic triad are indicated in blue. (E) *glit-1* gene structure with positions of the *glit-1(gt1981)* point mutation, the *glit-1(gk384527)* splice site mutation and the *glit-1(ok237)* deletion. For the point mutations the nucleotide and amino acid changes are indicated in brackets. The *ok237* deletion spans 5′UTR and first exon of *glit-1* and 5′UTR and first exons of *dnj-14* (DNaJ domain (prokaryotic heat shock protein)) and is indicated with a red bar. Also, *glit-1* is located in an operon (grey bar) with the ribosomal protein *rpl-25.1* (Ribosomal Protein, Large subunit).

### Loss-of-function mutations in the neuroligin-like *glit-1* cause 6-OHDA hypersensitivity

Combining whole-genome sequencing and single nucleotide polymorphism mapping [14,15], we determined that *gt1981* animals carry an amino acid-changing mutation in *glit-1 (<u>gli</u>o<u>t</u>actin (Drosophila neuroligin-like) homologue)* (Fig 1E). The *glit-1(gt1981)* 6-OHDA hypersensitivity phenotype was confirmed by the *glit-1(gk384527)* splice-site mutation and *glit-1(ok237)* promoter deletion (Fig 1C, E). The *glit-1(gk384527)* allele did not complement *gt1981; glit-1(gk384527)/gt1981* transheterozygous mutants were still hypersensitive to 6-OHDA (S1B Fig). To determine if the *glit-1(gt1981)* mutation behaves in a dominant or in a recessive fashion, *glit- 1(gt1981)*/*glit-1*(*gt1981)* homozygous mutants were crossed to wild-type (+) animals. We found that the *glit-1(gt1981)*/+ heterozygous F1 progeny did not exhibit increased 6-OHDA-induced neurodegeneration (S1C Fig). We conclude that loss-of-function of the neuroligin-like gene *glit-1* causes increased degeneration of dopaminergic neurons after exposure to 6-OHDA.

Neuroligins are postsynaptic cell adhesion proteins required for the correct function of synapses (for review [16]). The neuroligin protein family is defined by an extracellular carboxyesterase-like domain that is enzymatically inactive [16]. The neuroligin-like GLIT-1 is predicted to contain this conserved extracellular carboxyesterase-like domain, as well as a single-pass transmembrane domain and a short intracellular segment (S2 Fig). Phylogenetically, GLIT-1 groups with invertebrate esterases and its *Drosophila* homologue Gliotactin (S3 Fig). In the predicted carboxyesterase-like domain of GLIT-1, two aspartates and an arginine replace the characteristic catalytic triad of acetylcholinesterases (Fig 1D, S4 Fig, S5B-D Fig). Furthermore, the *glit-1(gt1981)* mutation which causes 6-OHDA sensitivity results in a proline to leucine amino acid substitution in the GLIT-1 carboxyesterase-like domain (Fig 1D, S5A Fig). This proline is likely important for protein function as it is conserved in neuroligins and acetylcholinesterases alike (S5A Fig). In summary, the *gt1981* mutation affects a conserved residue in GLIT-1, a previously uncharacterised *C. elegans* neuroligin-like protein.

Inhibition of neuronal 6-OHDA uptake by mutation of the *C. elegans* <u>d</u>op<u>a</u>mine <u>t</u>ransporter DAT-1 prevents 6-OHDA-induced dopaminergic neurodegeneration in wild-type animals [10]. We thus tested if DAT-1 is also required for 6-OHDA hypersensitivity conferred by *glit-1* mutation and found that this is the case as *glit-1;dat-1* double mutants were resistant to 6-OHDA (S1D Fig).

### *glit-1* is expressed in pharynx, intestine and several cells in the head, including dopaminergic neurons

To determine where GLIT-1 is expressed, we generated GFP reporter constructs. A P*glit-1∷gfp* transcriptional construct revealed that GLIT-1 is expressed in pharynx, intestine, and several cells in the head including dopaminergic neurons (S6F-H Fig). N-terminal GFP tagging of GLIT-1 protein confirmed this expression pattern (Fig 2A, B and S6A-E Fig). GFP∷GLIT-1 expression could be clearly detected in dopaminergic neurons in L4 stage larvae and adult animals, but such expression could not be ascertained at earlier developmental stages, possibly due to low levels of GLIT-1 expression (Fig 2C-F, S6C-E Fig, S1 Movie). We note that N-terminal GFP-tagging appears to render GLIT-1 non-functional, likely by compromising the N-terminal membrane targeting sequence: GFP∷GLIT-1 localised to the cytoplasm and not the plasma membrane. Accordingly, the construct did not rescue the 6-OHDA hypersensitivity of *glit-1(gt1981)* mutants (S7A Fig). Furthermore, a targeted chromosomal GFP insertion at the GLIT-1 C-terminus did not reveal any expression. In summary, transcriptional and translational reporter constructs show that GLIT-1 is expressed in pharynx, intestine and several cells in the head, including dopaminergic neurons.

**Fig 2.**
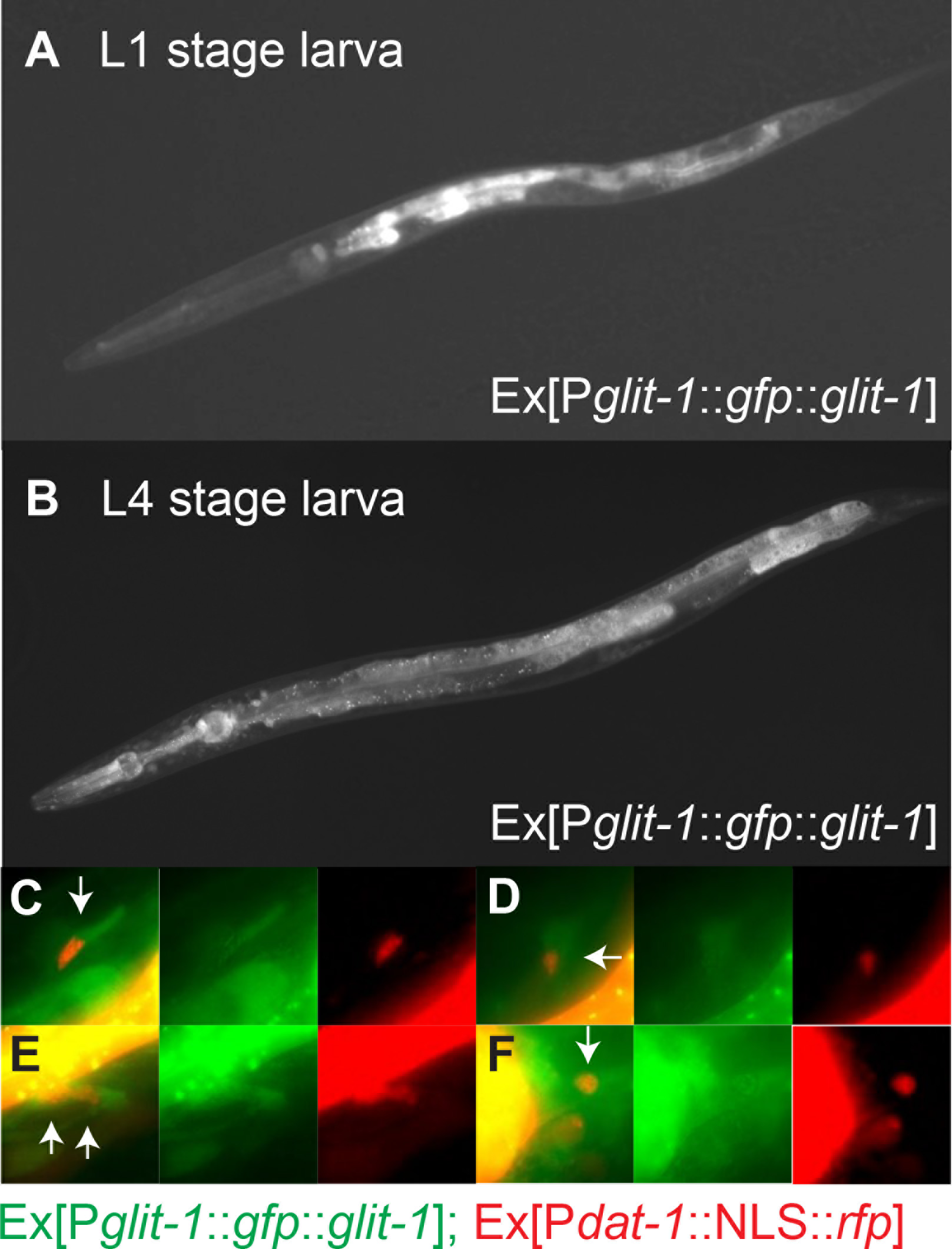
*glit-1* is expressed in pharynx, intestine and several cells in the head including dopaminergic neurons. (A) L1 stage larva expressing *Ex[Pglit-1∷gfp∷glit-1*]. (B) L4 stage larva expressing *Ex[Pglit-1∷gfp∷glit-1*]. This image was generated from three individual images in ImageJ using the Stitching plugin [60]. (C) – (F) Close up of dopaminergic neurons of same L4 stage larva. The green channel shows expression of *Ex[Pglit-1∷gfp∷glit-1*]. The red channel shows expression of *Is[Pdat-1∷NLS∷rfp;Pttx- 3∷mCherry*] for labelling of dopaminergic neuron nuclei, as well as the pharynx muscle marker *Ex[Pmyo-2∷mCherry*] and the body muscle marker *Ex[Pmyo-3∷mCherry*] that were used for injections.

We next aimed to determine in which tissues GLIT-1 acts to protect against toxicity conferred by 6-OHDA. An integrated, single copy construct driving *glit-1* expression under its own promoter almost completely prevented dopaminergic neuron loss (Fig 3A). In contrast, single-copy expression of *glit-1* in dopaminergic neurons, the pharynx and the intestine, respectively, did not alleviate 6-OHDA-induced neurodegeneration of *glit-1* mutants (Fig 3B, C). We conclude that GLIT-1 likely acts in several tissues to protect against dopaminergic neurodegeneration after 6-OHDA exposure.

**Fig 3.**
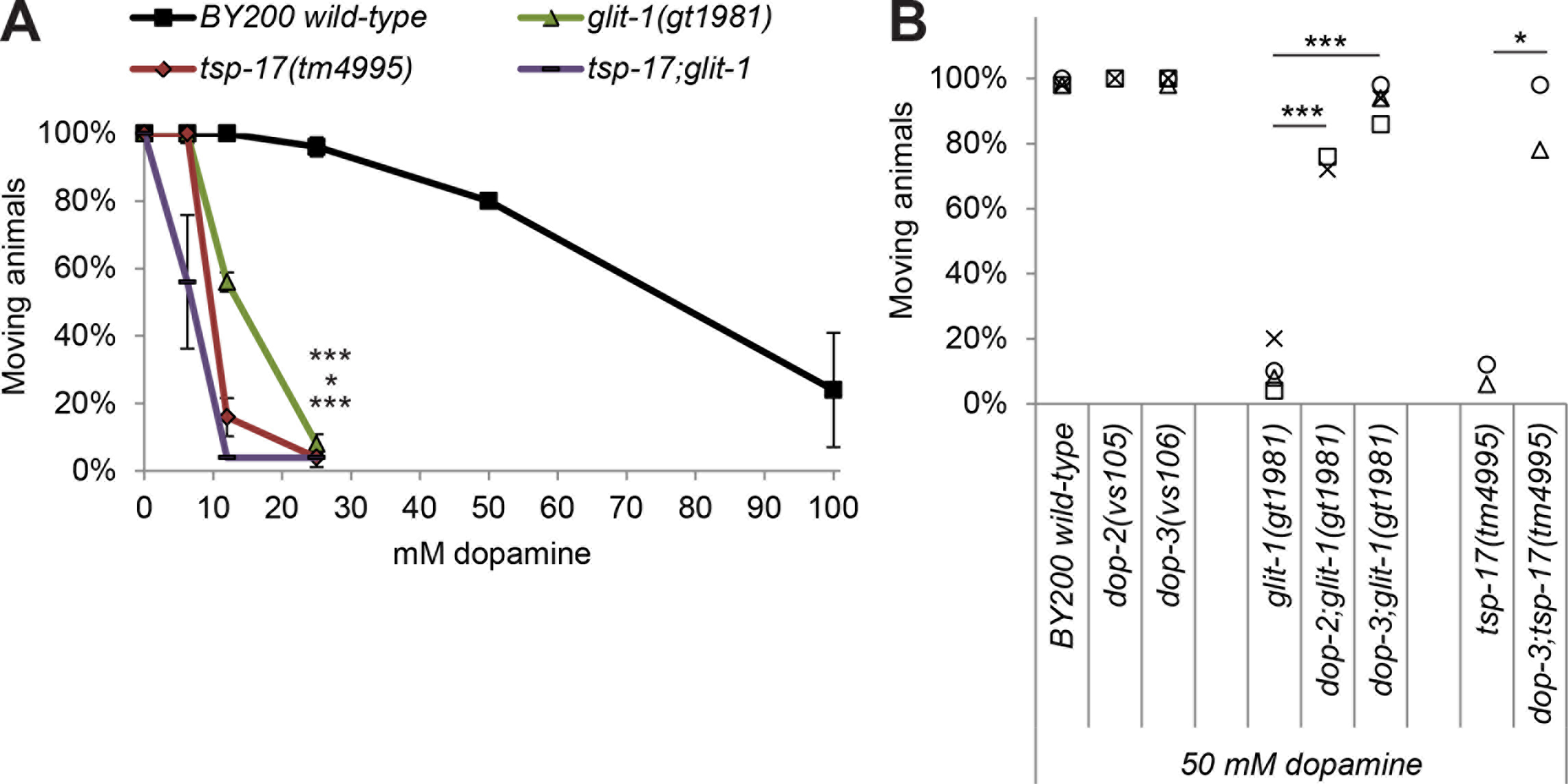
Single copy expression of *glit-1* alleviates neurodegeneration in *glit-1* mutants and *glit-1* likely functions in a pathway with the <u>t</u>etra<u>sp</u>anin *tsp-17*. (A) Effect of a single-copy *Is[Pglit-1∷glit-1*] construct on dopaminergic neurodegeneration after treatment with 10 mM 6-OHDA. Error bars = SEM of 2 biological replicates, each with 100-105 animals per strain. Total number of animals per condition n = 200-205 (****p<0.0001, **p<0.01; G-Test). (B) Effect of single-copy *Is[Pdat-1∷glit-1*] and Is[p*pha-4∷glit-1*] constructs on dopaminergic neurodegeneration after treatment with 10 mM 6-OHDA. Error bars = SEM of 2 biological replicates, each with 100-105 animals per strain. Total number of animals per condition n = 200-205 (n.s. p>.05; G-Test). (C) Effect of a single-copy Is[Pelt-2∷glit-1] construct on dopaminergic neurodegeneration after treatment with 10 mM 6-OHDA. Error bars = SEM of 2 biological replicates, each with 95-100 animals per strain. Total number of animals per condition n = 195-200 (n.s. p>0.05,; G-Test). (D) Dopaminergic head neurons in wild-type and *glit-1* and *tsp-17* single and double mutants 24, 48 and 72 hours after treatment with 0.75 mM 6-OHDA and 72 hours after control treatment with ascorbic acid only (’72 h Ctr’). Error bars = SEM of 2-3 biological replicates, each with 60-115 animals per strain. Total number of animals per condition n = 180-350 (*p<0.05, n.s. p>0.05; G-Test). The ‘72 h Ctr’ was conducted twice with 30 animals per strain, resulting in a total n = 60.

To test if GLIT-1 overexpression protects wild-type animals against dopaminergic neuron loss after 6- OHDA exposure, we analysed two independent transgenic lines carrying *Pglit-1∷glit-1* extrachromosomal arrays. The extent of neurodegeneration in these transgenic animals was comparable to that of wild-type animals, suggesting that overexpression of *glit-1* does not confer protection against 6-OHDA (S7B Fig).

### GLIT-1 acts in the same genetic pathway as TSP-17

Previously, we have shown that mutation of the *C. elegans* <u>t</u>etra<u>sp</u>anin *tsp-17* renders dopaminergic neurons hypersensitive to 6-OHDA [13]. To determine if GLIT-1 and TSP-17 act in the same genetic pathway, we generated the *glit-1(gt1981);tsp-17(tm4995)* double mutant and found that it exhibits a similar extent of 6-OHDA-induced dopaminergic neurodegeneration as the *glit-1* and *tsp-17* single mutants (Fig 3D). Hence, GLIT-1 and TSP-17 are likely to function in the same genetic pathway to protect from dopaminergic neuron loss after exposure to 6-OHDA.

We used qRT-PCR to determine if expression of *tsp-17* and *glit-1* is induced after 6-OHDA treatment. However, we did not find altered *tsp-17* and *glit-1* transcript levels in wild-type and *tsp-17* and *glit-1* mutant larvae (S7C-F Fig), suggesting that *tsp-17* and *glit-1* provide constitutive protection against oxidative stress. This hypothesis is consistent with the fact that we did not determine a protective effect of *glit-1* overexpression (S7B Fig).

### GLIT-1 does not appear to affect intestinal permeability and acts independently of NLG-1 and NRX-1

The *glit-1 Drosophila* homologue Gliotactin is localised at tricellular junctions and necessary for the development of septate junctions, an invertebrate analogue of tight junctions in epithelial cells [17,18]. To address if *glit-1* mutation might compromise the epithelial barrier function of the *C*. *elegans* intestine, we tested if a blue dye (‘Smurf’ assay) or fluorescein leak into the body cavity after ingestion [19,20]. We did not find an increased intestinal permeability in *glit-1* mutants as compared to wild-type animals (S8 Fig and S9 Fig). We note that the molecular weight of fluorescein (Mw = 376.2 g/mol), the smaller one of the two dyes, is still slightly higher than the molecular weight of 6-OHDA (Mw = 205.64 g/mol). Thus, we cannot fully exclude the possibility of increased organismal 6-OHDA uptake in *glit-1* mutants, albeit this appears unlikely.

*C. elegans* possesses two neuroligin genes: The neuroligin-like *glit-1*, which forms the centre of this study, and the neuroligin *nlg-1*. NLG-1 has been shown to mediate oxidative stress protection [21,22] and is transcriptionally induced by the oxidative stress ‘master regulator’ SKN-1 (<u>sk</u>i<u>n</u>head)/Nrf [22]. To test for genetic interactions between *glit-1* and *nlg-1*, we scored dopaminergic neurodegeneration in single and double mutants after treating animals with 0.75 mM 6-OHDA, a concentration that leads to a loss of approximately 50% dopaminergic head neuron in *glit-1*-mutants (S10A Fig) and thus allows for assessing increased or decreased neurodegeneration. We found that *nlg-1* mutation does not influence 6-OHDA-induced dopaminergic neurodegeneration in wild-type or *glit-1* mutant animals (S10B, C Fig). As neuroligins generally modulate signalling by contacting pre-synaptic neurexins [16], we further asked if *glit-1* genetically interacts with the *C. elegans* neurexin *nrx-1*. We tested two different neurexin deletions; *nrx-1(wy778)* affecting long and short isoform of the gene, and *nrx-1(ok1649)* affecting only long neurexin isoforms. We found that while *nrx-1(wy778)* mutation does not influence 6-OHDA-induced neurodegeneration, *nrx-1(ok1649)* mutation slightly alleviated dopaminergic neuron loss in *glit-1* mutants and wild-type animals alike (S10B-D Fig). These results indicate that there are no specific genetic interactions between the neuroligin-like *glit-1* and the neuroligin *nlg-1* or between *glit-1* and neurexin that affect 6-OHDA-induced dopaminergic neuron loss.

### *glit-1* mutation does not appear to affect dopamine signalling

As *glit-1* and *tsp-17* are both expressed in dopaminergic neurons, we aimed to determine if the hypersensitivity of *glit-1* mutants to 6-OHDA is altered by mutations affecting dopamine metabolism (simplified cartoon in S11F Fig., for review [23]). However, we did not find that dopamine receptor mutations specifically alter 6-OHDA-induced dopaminergic neurodegeneration in the *glit-1* mutant background (S11A, D, E Fig, S1A Text). We further found that some mutants defective in dopamine synthesis can lead to increased 6-OHDA-induced neurodegeneration specifically in *glit-1* mutants (S11B – E Fig). At present, we cannot explain the partly contradictory effects we found (S1B Text). To assess in more detail if the signalling of dopaminergic neurons is altered by *glit-1* and *tsp-17* mutation, we tested several behaviours that are mediated by dopamine. We firstly found that dopamine signalling appears to be functional in the mutants [13] (S12A Fig) by analysing the basal slowing response, the dopamine-mediated slowing of *C. elegans* when encountering a lawn with bacterial food [24]. Secondly, we found that *glit-1* mutants (S12B Fig), unlike *tsp-17* mutants [13], do not exhibit premature swimming-induced paralysis, a behaviour indicative of increased extracellular dopamine levels [25]. Thirdly, we tested dopamine-induced paralysis: Overstimulation of *C. elegans* with exogenous dopamine eventually leads to animal paralysis [26]. We found that *glit-1* and *tsp-17* mutants stopped moving at much lower dopamine concentrations than wild-type animals (Fig 4A). This increased sensitivity to exogenous dopamine depends on dopamine signalling as it was rescued by mutation of the dopamine receptors *dop-2* and *dop-3* (Fig 4B). We conclude that *glit-1* mutants, in contrast to *tsp-17* mutants, do not exhibit signs of altered intrinsic dopamine signalling. However, both *tsp-17* and *glit-1* mutants exhibit increased dopamine-induced paralysis.

**Fig 4.**
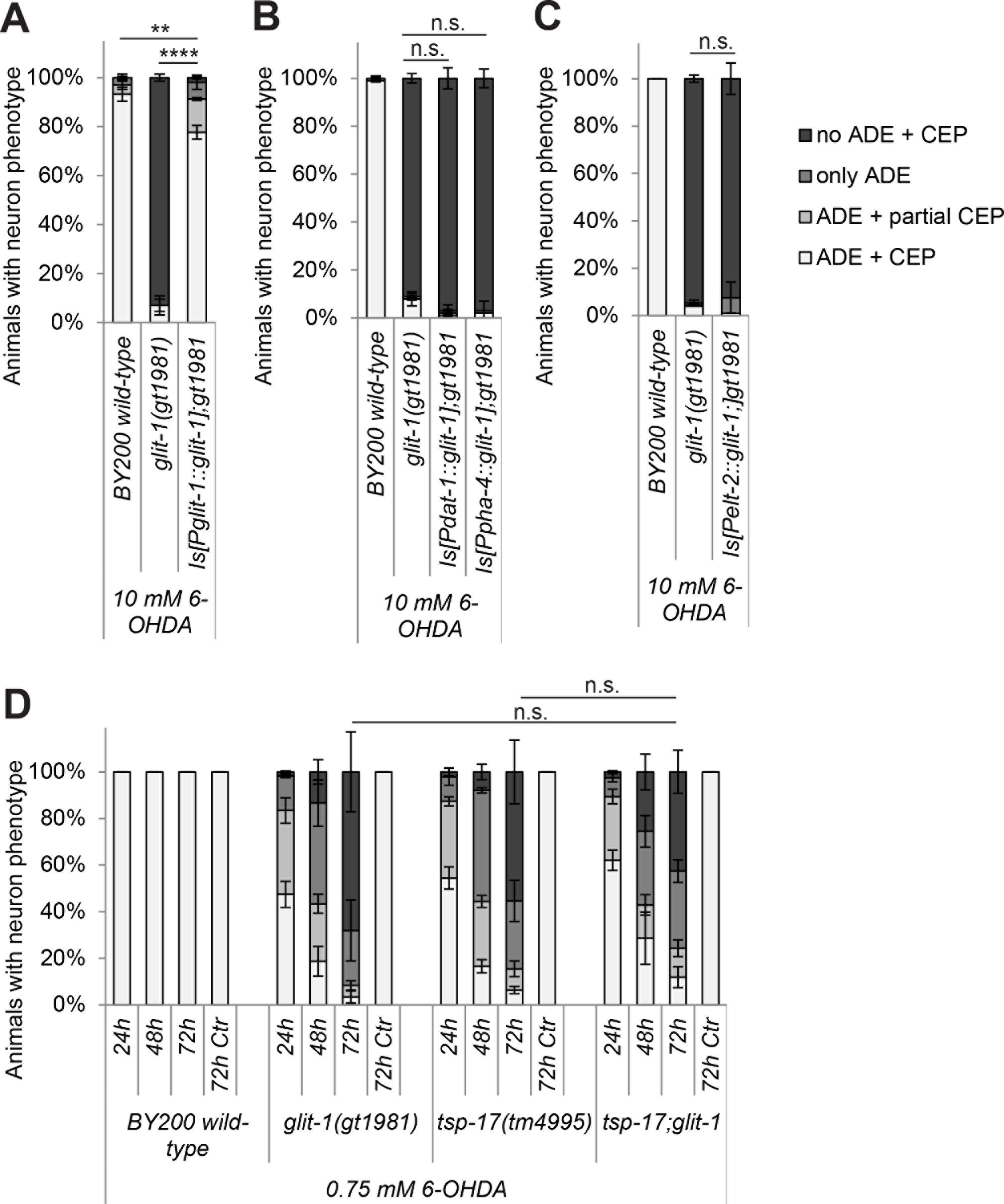
*glit-1* and *tsp-17* exhibit increased dopamine-induced paralysis. (A) Percentage of moving young adults on plates with indicated concentrations of dopamine. Error bars = StDev of 2 technical replicates, each with 25 animals per strain and condition. Total number of animals per condition n = 50 (***p<0.001, *p<0.05, n.s. p>0.05; two-tailed t-test comparing wild- type and mutant animal data at 25 mM dopamine). (B) Percentage of moving young adults on plates containing 50 mM dopamine. Indicated with grey symbols are the 2-4 biological replicates, each performed with 50 animals per strain. Total number of animals per condition n = 100-200 (***p<0.001, *p<0.05; two-tailed t-test).

### *glit-1* and *tsp-17* mutants are sensitive to oxidative stress at an organismal level

As oxidative stress elicited by 6-OHDA led to increased dopaminergic neurodegeneration in *glit-1* and *tsp-17* mutant animals, we aimed to determine if these mutants are also hypersensitive to oxidative stress at an organismal level. To this end we exposed animals for 1 hour to paraquat, an oxidative stress-inducing herbicide linked to the development of Parkinson’s disease [3]. Paraquat, unlike 6-OHDA, is not known to be specifically taken up into dopaminergic neurons. We found that after incubation with different concentrations of paraquat, *glit-1* and *tsp-17* mutants exhibit a very similar developmental delay compared to wild-type animals (Fig 5A). To follow up on this result, we also analysed the second *glit-1* allele *glit-1(gk384527)*, as well as the *tsp-17(tm4995);glit-1(gt1981)* double mutant after exposure to very low concentrations of paraquat. We found that *tsp-17* and *glit-1* single and double mutants exhibit an almost identical developmental delay phenotype (Fig 5B). Furthermore, *glit-1* and *tsp-17* mutants also developed much slower than wild-type animals after incubation with hydrogen peroxide (H2O2), another oxidative stress-inducing compound (Fig 5C). In summary, *glit-1* and *tsp-17* mutants are hypersensitive to oxidative stress at the organismal level and likely act in the same pathway.

**Fig 5.**
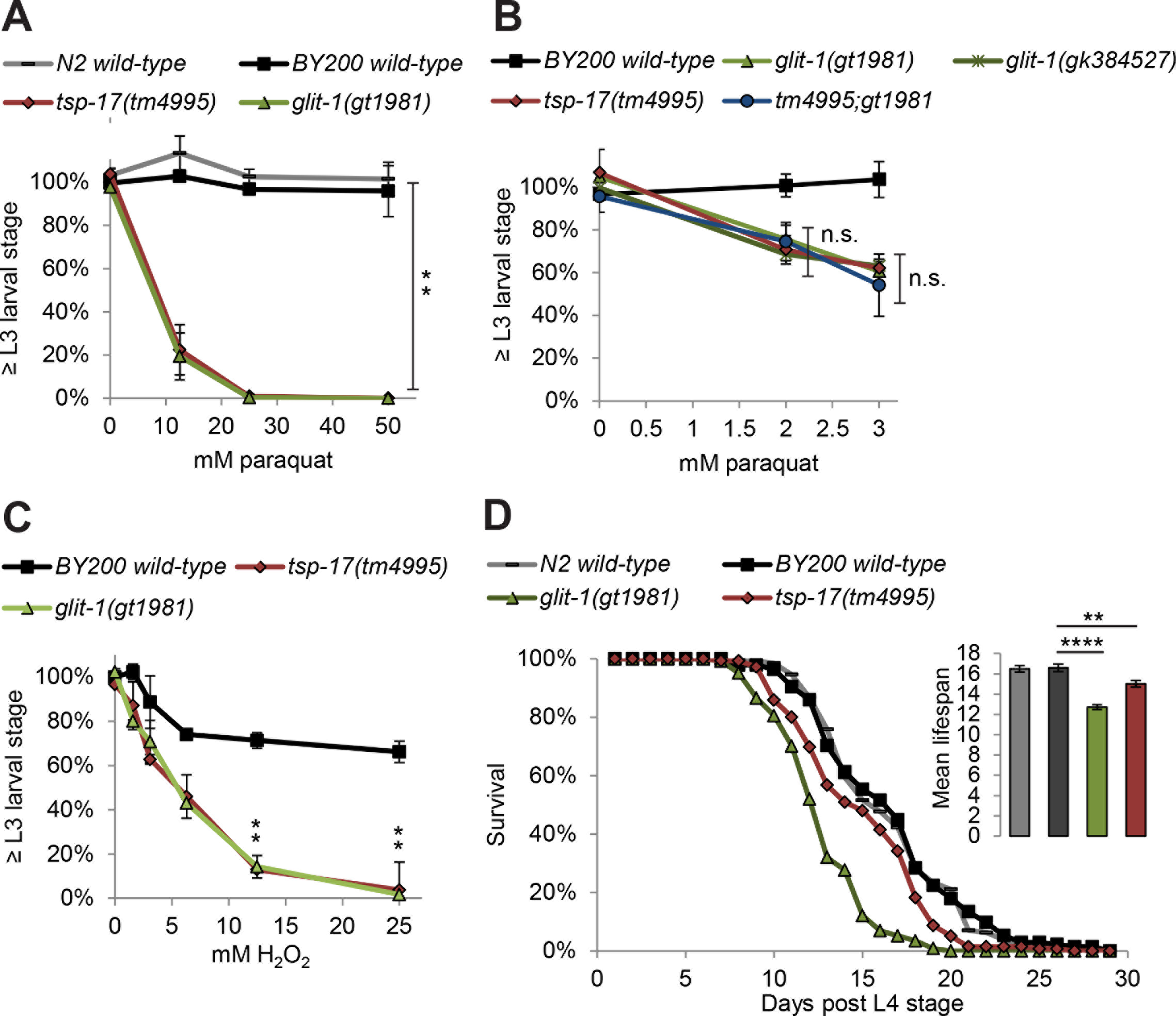
*glit-1* and *tsp-17* are hypersensitive to oxidative stress at the organismal level. (A) Percentage of animals that developed to L3 stage 24 hours after 1 hour incubation with indicated concentration of paraquat. Error bars = SEM of 3 biological replicates, each with 85-365 animals per strain and concentration. Total number of animals per condition n = 350-650 (*p<0.05; two-tailed t- test comparing data of BY200 wild-type and mutant animal data at 50 mM paraquat). (B) Percentage of animals that developed to L3 stage 24 hours after 1 hour incubation with 50 mM paraquat. Error bars = SEM of 3 biological replicates, each with 85-380 animals per strain and concentration. Total number of animals per condition n = 340-830 (n.s. p>0.05; two-tailed t-test comparing the *tsp- 17(tm4995)* and *glit-1(gt1981)* single mutants to the *tsp-17(tm4995);glit-1(gt1981)* double mutant at 2 and 3 mM paraquat). (C) Percentage of animals that developed to L3 stage or beyond 48 hours after 1 hour incubation with indicated concentration of H_2_O_2_. Error bars = SEM of 2 biological replicates, each with 165-547 animals per strain and concentration. Total number of animals per condition n = 311-942 (*p<0.05; two-tailed t-test comparing data of BY200 wild-type and mutant animal data at 12.5 and 25 mM H_2_O_2_). (D) Lifespan data for first biological replicate including 97-130 animals per strain. The inset shows the mean lifespan with the error bars depicting the standard error (****p<0.0001, **p<0.01; Bonferroni-corrected; Log-Rank Test).

Increased oxidative stress has been implicated with premature aging (for review [27]), thus we asked if the increased oxidative stress sensitivity exhibited by *glit-1* and *tsp-17* mutants is linked to a shorter lifespan. We found that *glit-1* and *tsp-17* mutants die earlier than wild-type animals (Fig 5D, S13 Fig), indicating that the overall fitness of the mutants is decreased.

### Oxidative stress reporters are hyper-induced in *glit-1* and *tsp-17* mutants

We aimed to determine if *glit-1* and *tsp-17* mutants exhibit an altered oxidative stress response. Using qRT-PCR, we measured transcript levels of the oxidative stress reporter *gst-4 (glutathione S-transferase), gcs-1 (gamma glutamylcysteine synthetase)* and *gst-1 (glutathione S-transferase)* in wild-type and mutant L1 stage larvae after 1 hour of incubation with 10 mM 6-OHDA in comparison to mock treatment. GST-1 was previously shown to mildly inhibit the drug-induced degeneration of *C. elegans* dopaminergic neurons [28]. We found that *gst-4* transcripts were approximately 1.5-fold upregulated in wild-type animals after 6-OHDA treatment in comparison to approximately 5-fold in *tsp-17* and *glit-1* mutants (Fig 6A and S15A Fig). Furthermore, *gcs-1* and *gst-1* transcripts were not induced by 6-OHDA in wild-type larvae, but an approximately 2-3-fold induction occurred in *tsp-17* and *glit-1* mutants (Fig 6B, C and S15B, C Fig). Comparing the basal level of *gcs-1* and *gst-1* and *gst-4* expression in wild-type and *tsp-17* and *glit-1* mutants, we found that there is no difference in *gcs-1* and *gst-1* expression, while *gst-4* appeared upregulated in *tsp-17* and *glit-1* mutants. (S14 and S15D- F Fig). In addition, we measured reporter expression after exposure to paraquat and found a modest level of hyperinduction of the reporters in *tsp-17* and *glit-1* mutants (S16 Fig). In summary, we found increased levels of oxidative stress signalling in *glit-1* and *tsp-17* mutants, indicating that the two genes protect from various oxidative-stress inducing compounds.

**Fig 6.**
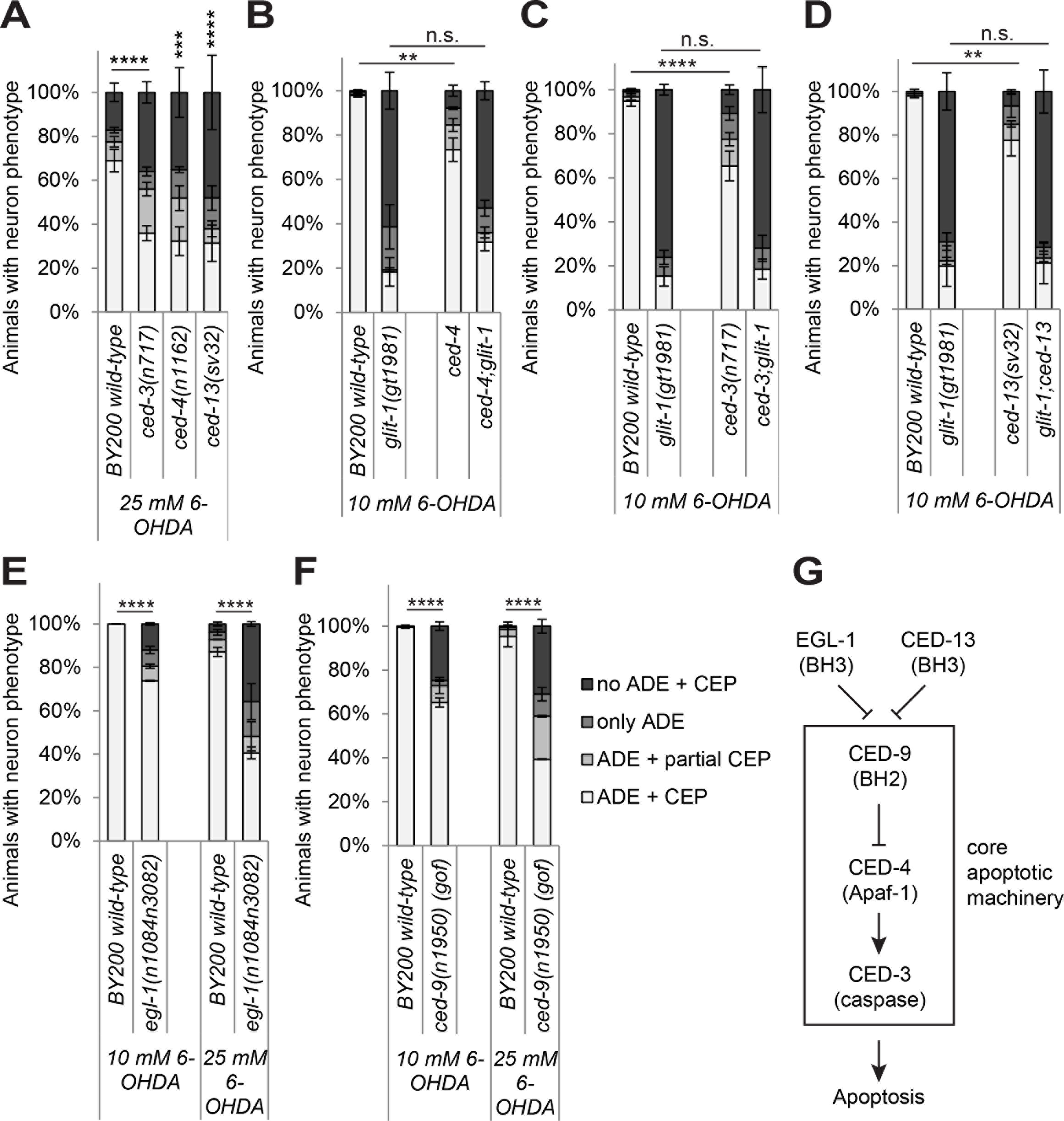
Expression of oxidative stress reporters is increased in *glit-1* and *tsp-17* mutants and hyperactivation of *skn-1* alleviates 6-OHDA-induced neurodegeneration. (A) *gst-4* (B) *gcs-1* and (C) *gst-1* mRNA levels in wild-type and mutant L1 stage larvae after 1 hour treatment with 10 mM 6-OHDA. (A)-(C) The data are normalised to the control gene *Y45F10D.4*. The average and the respective values for 5-6 biological replicates (biorep a-f) are indicated. Error bars = SEM of 5-6 biological replicates (**p<0.01, *p<0.05; two-tailed t-test). (D) Effect of *skn-1 gain-of- function (gof)* mutations *skn-1(lax120)* and *wdr-23(tm1817)* on dopaminergic neurodegeneration after treatment with 0.75 mM 6-OHDA. Error bars = SEM of 3-4 biological replicates, each with 40120 animals per strain and concentration. Total number of animals per strain n = 180-430 (****p<0.0001, **p<0.01; G-Test). (E) Effect of *skn-1 gain-of-function (gof)* mutations *skn-1(lax120)* and *wdr-23(tm1817)* on dopaminergic neurodegeneration after treatment with 50 mM 6-OHDA. Error bars = SEM of 3-6 biological replicates, each with 70-115 animals per strain and concentration. Total number of animals per condition = 290-430 (****p<0.0001; G-Test). (F) Effect of *skn-1 loss-of- function (lof)* allele *zu67* on dopaminergic neurodegeneration after treatment with 10 mM 6-OHDA. Error bars = SEM of 3 biological replicates, each with 30-105 animals per strain. Total number of animals per strain n = 170-270 (n.s. p>0.05; G Test). Experiment was conducted with unstarved, filtered L1 larvae since an insufficient amount of *skn-1(zu67)* mutants could be gained in liquid culture. Unstarved animals show a higher degree of degeneration.

Both *gst-4* and *gcs-1* are targets of SKN-1 [29,30], an orthologue of the mammalian Nrf (nuclear factor E2-related factor) transcription factor family (for review [31]) which promotes oxidative stress resistance and longevity in *C. elegans* [32]. Thus, we aimed to determine if SKN-1 is involved in the protection against 6-OHDA-induced dopaminergic neurodegeneration. In *C. elegans*, SKN-1 abundance is negatively regulated by the CUL-4/DDB-1 ubiquitin ligase substrate targeting protein WDR-23 (<u>WD</u> <u>R</u>epeat protein) [33]. We found that hyperactivation of SKN-1 activity – conferred by a *gain-of-function (gof)* mutation in *skn-1* or by loss of the SKN-1 repressor WDR-23- led to reduced dopaminergic neurodegeneration in *glit-1* mutants (Fig 6D) and wild-type animals (Fig 6E). However, a *skn-1(zu67) loss-of-function* mutant did not exhibit increased dopaminergic neuron loss (Fig 6F). Thus, SKN-1 hyperactivation generally improves the defence against 6-OHDA-induced neurodegeneration, but it is unlikely that SKN-1 malfunction underlies 6-OHDA hypersensitivity observed in *glit-1* and *tsp-17* mutants.

We tested additional *C. elegans* stress response pathways for their involvement in 6-OHDA protection. However, we found that mutation of the DAF-2/DAF-16 insulin/insulin-like growth factor signalling pathway (for review [34]), the JNK-1 and KGB-1 JNK (<u>J</u>un <u>N</u>-terminal <u>k</u>inase) pathways and the PMK-1 and PMK-3 p38 MAPK (<u>m</u>itogen-<u>a</u>ctivated <u>p</u>rotein <u>k</u>inase) pathways (for review [35]) did not overtly alter 6-OHDA-induced dopaminergic neuron loss (S17 Fig, S18 Fig). In addition, we tested if 6-OHDA induces the expression of reporters for the endoplasmic reticulum unfolded protein response, *hsp-4*, and the mitochondrial unfolded protein response, *hsp-6*. While *hsp-4* transcript levels were slightly elevated after 6-OHDA, we did not detect a change in *hsp-6* transcript levels (S19C-F Fig). Furthermore, we did not find a difference in *hsp-4* and *hsp-6* transcript levels between wild-type, and *tsp-17* and *glit-1* mutant larvae (S19C, D, G, H Fig). Thus, of the stress pathways we tested, only hyperactivation the *skn-1* pathway leads to detectable protection from neurotoxicity conferred by 6-OHDA.

### Mutations affecting the apoptosis pathway sensitise dopaminergic neurons to 6-OHDA

It is not known which cell death pathway mediates 6-OHDA-induced neurodegeneration, so we revisited the role of the canonical apoptosis pathway. The conserved core apoptosis pathway is required for the vast majority of *C. elegans* apoptotic cell deaths and comprised of proapoptotic EGL-1 BH3-only domain protein, the Apaf1-like protein CED-4 and the CED-3 caspase. The apoptosis pathway also includes the Bcl2-like protein CED-9, which protects from apoptosis. CED-13 in contrast solely contributes to apoptosis induced by DNA damaging agents such as ionising irradiation[36] (simplified overview in Fig 7G, for review [37]). In addition to mediating apoptosis, CED-3, CED-4 and CED-13 were also reported to be part of a hormetic response mediating lifespan extension upon limited mitochondrial oxidative stress [38]. We found that compared to wild-type animals, *egl-1, ced-13, ced-4* and *ced-3* loss-of-function (lof) mutants and the *ced-9* gain-of-function (gof) mutant exhibit increased 6-OHDA-induced dopaminergic neurodegeneration (Fig 7A-F). Thus, the apoptosis pathway appears to protect dopaminergic neurons after 6-OHDA exposure. In contrast, we found that increased neurodegeneration in *glit-1* mutants is not further enhanced by *ced-3, ced-4* and ced- 13 mutations (Fig 7B-D, S20 Fig). It is conceivable that the increased neurodegeneration conferred by apoptosis gene mutations becomes undetectable in the *glit-1* mutant background. Alternatively, it is possible that those apoptosis genes and *glit-1* act in the same genetic pathway to protect dopaminergic neurons. In summary, our results show that the apoptosis pathway helps to prevent 6- OHDA-induced dopaminergic neurodegeneration.

**Fig 7.**
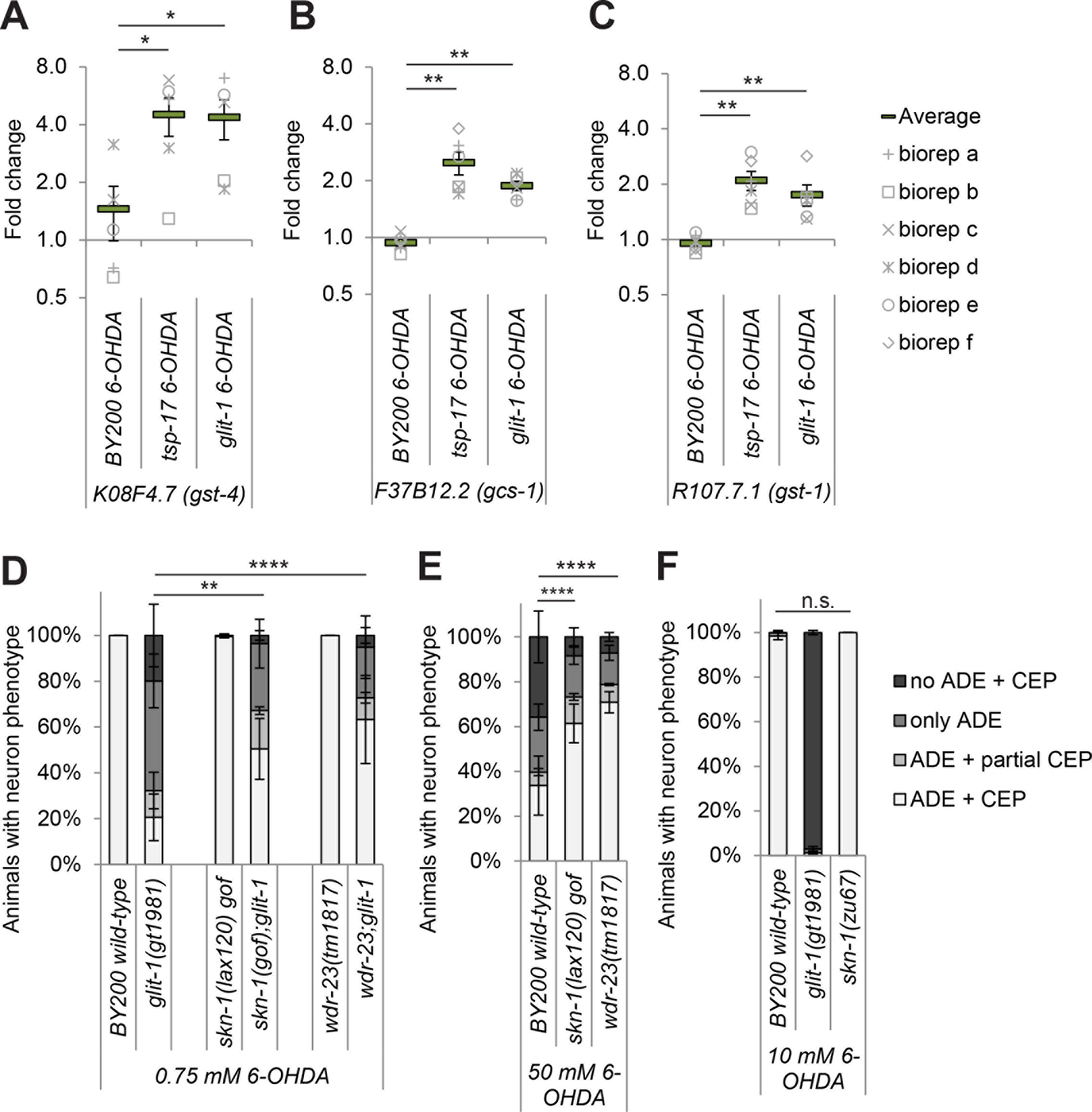
Mutations in the canonical apoptosis pathway sensitise to 6-OHDA-induced dopaminergic neurodegeneration. (A) Effect of apoptosis/mtROS pathway mutations on dopaminergic neurodegeneration after treatment with 25 mM 6-OHDA. Error bars = SEM of 3 biological replicates, each with 70-120 animals per strain and concentration. Total number of animals per condition n = 280-320 (****p<0.0001, ***p<0.001, n.s. p>0.05; G-Test). (B) Effect of *ced-4*, (C) *ced-3* (D) *ced-13*, (E) *egl-1 loss-of-function* mutations, as well as the (F) *ced-9 gain-of-function (gof)* mutation on dopaminergic neurodegeneration after treatment with 10 mM 6-OHDA. Error bars = SEM of 2-5 biological replicates, each with 40-115 animals per strain and concentration. Total number of animals per condition n = 150-490 (****p<0.0001, **p<0.01, n.s. p>0.05; G-Test). (G) Simplified scheme of the apoptosis pathway (adapted from [61]).

### Mutation of the calcium-storing chaperone *crt-1* exacerbates 6-OHDA-induced dopaminergic neurodegeneration

Since dopaminergic neurons do not undergo apoptosis after 6-OHDA exposure, we aimed to determine if they die via a necrosis-like cell death. Mutation of the *C. elegans* calreticulin CRT-1, a calcium-binding and calcium-storing chaperone in the endoplasmic reticulum, prevents several necrosis-type cell deaths [39,40]. In contrast, we found that *crt-1* mutation led to a highly increased loss of dopaminergic neurons after exposure to 6-OHDA in an otherwise wild-type background (Fig 8A-C). We further found that the *crt-1;glit-1* and the *crt-1;tsp-17* double mutants are synthetic lethal. We used a fluorescently labelled balancer chromosome to keep *crt-1* in a heterozygous state in a *glit-1* homozygous background. In the next generation, genetic segregation of the *crt-1* and the fluorescent balancer chromosome is expected. However, we did not find any non-fluorescent animals, confirming that *crt-1* homozygosity is lethal in the *glit-1* mutant background. We furthermore found that synthetic lethality occurs when *tsp-17(tm4995)* and *crt-1(ok948)* are combined. Although it is possible that this synthetic lethality is related to the function of CRT-1 in calcium-induced necrosis, synthetic lethality might also be caused by developmental roles of *crt-1*: *crt-1* mutation was shown to be synthetic lethal with *vab-10* and other genes required for mechanical resilience of the epidermis [41]. Using qRT-PCR, we also tested if *crt-1* expression is altered by 6-OHDA but we found *crt-1* transcript levels remain unchanged after 6-OHDA treatment in wild-type, and in *tsp-17* and *glit-1* mutant animals (Fig 8D, S21B Fig). *crt-1* expression also remains constant after paraquat treatment (S16 Fig). Furthermore, basal expression of *crt-1* is not altered in *tsp-17* or *glit-1* mutants as compared to wild-type (Fig 8E, S21C Fig). To follow up on a more general role of calcium homeostasis in 6-OHDA-induced dopaminergic neurodegeneration, we also analysed *itr-1 ([nositol triphosphoate receptor)* and *cnx-1 (<u>c</u>al<u>n</u>e<u>x</u>in)* mutants. ITR-1 and CNX-1 were proposed to play similar roles as CRT-1 in the calcium-induced necrotic cell death of *C. elegans* PLM (<u>p</u>osterior <u>l</u>ateral <u>m</u>icrotubule cells) neurons [39]. However, we found that unlike *crt-1* mutation, *itr-1* and *cnx-*1 mutations do not alter the extent of dopaminergic neuron loss after 6-OHDA exposure (S21A Fig). In summary, we report that the calreticulin orthologue CRT-1 is required for protecting against 6- OHDA-mediated dopaminergic neurodegeneration.

**Fig 8.**
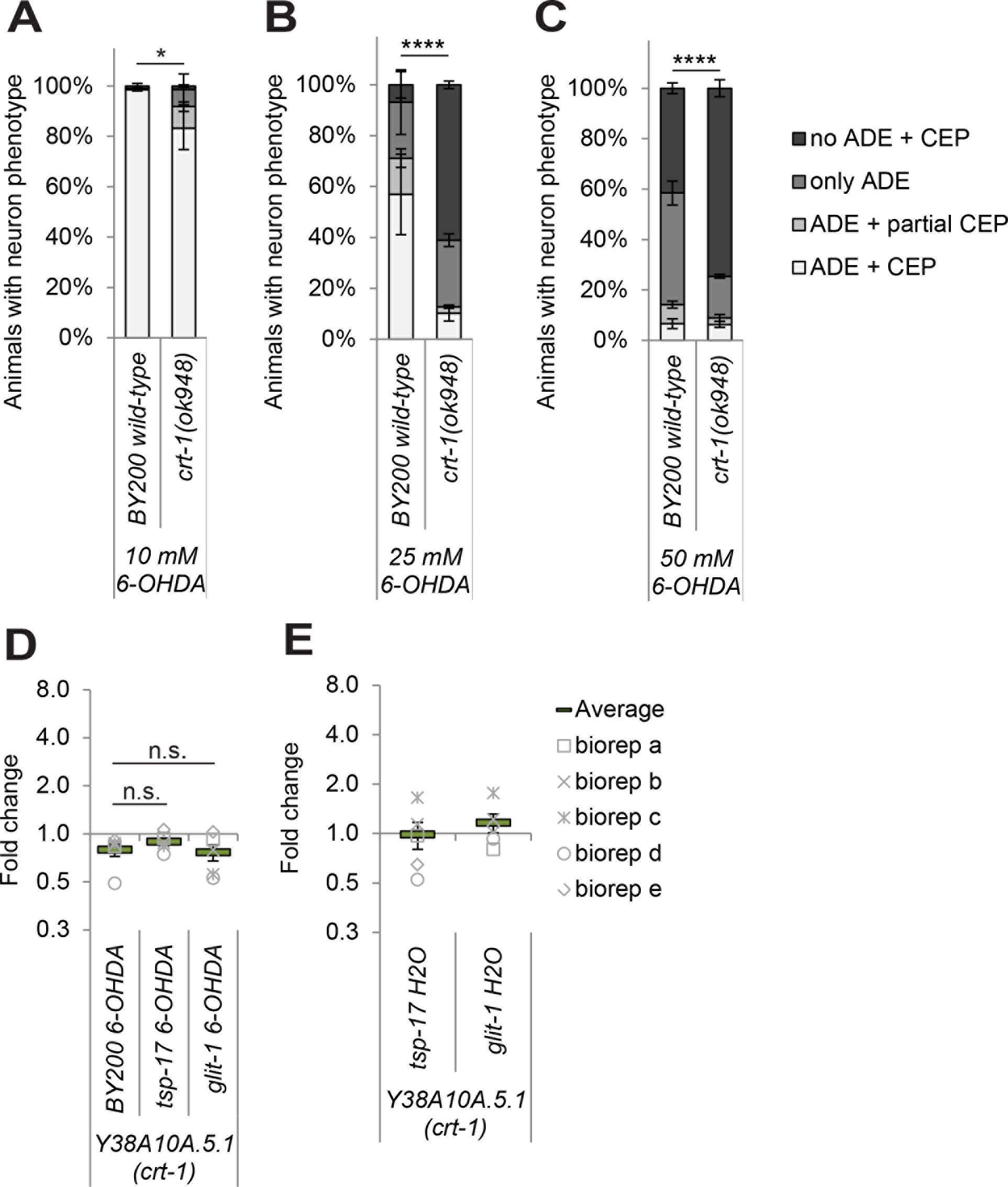
Mutation of the calcium chaperone *crt-1* sensitises to 6-OHDA-induced dopaminergic neurodegeneration. (A) Effect of mutations in *crt-1* mutation on dopaminergic neurodegeneration after treatment with 10 mM, (B) 25 mM and (C) 50 mM 6-OHDA. Error bars = SEM of 2-3 biological replicates with 90-130 animals per strain and concentration. Total number of animals per strain = 200-360 (****p<0.0001, *p<0.05; G-Test). (D) *crt-1* mRNA level analysis after 1h of 6-OHDA treatment in wild-type and mutant L1 stage larvae. (E) *crt-1* mRNA level analysis in mutant L1 stage larvae as compared to wild- type animals under control conditions (treatment with H_2_O instead of 6-OHDA). (D) and (E) The data are normalised to the control gene *Y45F10D.4*. The average and the respective values for 5 biological replicates (biorep a-e) are indicated. Error bars = SEM of 5 biological replicates (n.s. p>0.05; twotailed t-test).

## Discussion

We show that GLIT-1 protects *C. elegans* dopaminergic neurons from 6-OHDA-induced degeneration, likely in a similar way as the previously identified <u>t</u>etra<u>sp</u>anin TSP-17 [13]. We found that *glit-1* and *tsp-17* mutants are equally hypersensitive to 6-OHDA and paraquat, and that double mutants do not exhibit an enhanced phenotype. We previously showed that *tsp-17* mutants display phenotypes consistent with excessive dopaminergic signalling and we hypothesised that TSP-17 protects dopaminergic neurons by inhibiting the uptake of 6-OHDA into dopaminergic neurons [13]. In this study, we uncovered that both TSP-17 and GLIT-1 protect against oxidative stress at an organismal level. Consistent with increased oxidative stress levels in *glit-1* and *tsp-17* mutants, we find increased transcription of oxidative stress reporters and hypersensitivity after exposure to oxidative stress-inducing compounds. In addition, using a reverse genetics approach, we found that the calcium chaperone CRT-1, as well as canonical apoptosis pathway and CED-13, a protein previously implicated in DNA damage-induced apoptosis, are also required to protect dopaminergic neurons from 6-OHDA-mediated degeneration.

We are considering several possibilities as to how the GLIT-1 and TSP-17 transmembrane proteins might confer the various phenotypes we observe. *glit-1* and *tsp-17* are both expressed in dopaminergic neurons. However, in contrast to TSP-17, which also appears to partially function in dopaminergic neurons [13], GLIT-1 expression is likely required in multiple tissues to confer protection against 6-OHDA. Furthermore, *tsp-17* mutants, but not *glit-1* mutants display subtle defects in a behaviour mediated by dopamine, the swimming-induced paralysis (this study and [13]). Both the neuroligin-like *glit-1* mutant and the <u>t</u>etra<u>sp</u>anin *tsp-17* mutant exhibit increased dopamine-induced paralysis. This dopamine sensitivity phenotype could point towards an increased level of intrinsic dopamine signalling, possibly accounting for an increased level of 6-OHDA uptake via the DAT-1 transporter. It is also conceivable that the increased sensitivity to dopamine-induced paralysis and oxidative stress is due to a higher organismal drug uptake, which might be caused by decreased endothelial barrier function. The GLIT-1 homologue in *Drosophila*, Gliotactin, localises to tricellular junctions and is necessary for the development of septate junctions, an invertebrate analogue of epithelial tight junctions [17,18]. However, using dye leakage assays, we did not find indications for increased intestinal permeability in *glit-1* mutant animals.

We consider it likely that both *glit-1* and *tsp-17* protect constitutively against oxidative stress at an organismal level, as several oxidative stress reporters are hyper-induced in *glit-1* and *tsp-17* mutants. The specific sensitivity of dopaminergic neurons to 6-OHDA is likely due to the specific uptake of 6-OHDA via the dopamine transporter. In line with this, we never observed dopaminergic neurodegeneration in wild-type or mutant animals after treatment with paraquat and H_2_O_2_, even at doses at which animal development is severely delayed. Both TSP-17 and GLIT-1 are homologous to vertebrate <u>t</u>etra<u>sp</u>anins and neuroligin-like molecules, but do not have clear mammalian orthologues. As the genetic screen for factors that protect dopaminergic neurons from oxidative damage is far from saturation, we consider it likely that further studies like ours will help to uncover additional pathways that protect neurons from oxidative damage.

Our data clearly indicate that the apoptosis pathway protects dopaminergic neurons from 6-OHDA-induced degeneration. The increased neuronal loss in *egl-1 loss-of-function (lof), ced-9 gain-of-function, ced-3 (lof)* and *ced-4 (lof)* mutants could be ascribed to a defect in the core apoptosis pathway defined by these genes. However, the *ced-13 (lof)* mutant also displays higher dopaminergic neurodegeneration. CED-13 has previously been found to mildly affect radiation-induced germ cell apoptosis, but not to alter developmental apoptosis as mediated by the core apoptosis pathway [36]. In addition, the core apoptosis pathway and CED-13 were reported to form a protective, lifespan-prolonging pathway that senses mild oxidative stress (conferred by the partial inhibition of mitochondrial respiration) [38]. Remarkably, *glit-1* expression was increased under these oxidative stress conditions (supplementary information in [38]). It is still not known how apoptosis proteins sense oxidative stress [38], and it certainly remains to be established how they confer protection of dopaminergic neurons after exposure to 6-OHDA (our observations). There is mounting evidence that ‘apoptosis proteins’ might adopt functions that are not related to apoptosis. For instance, it was demonstrated that *ced-4* is required for DNA damage-induced cell cycle arrest [42] and we previously showed that CED-4 and CED-9 are expressed in both dying and surviving cells, consistent with roles beyond apoptosis [43].

We further found that the calreticulin CRT-1 protects against 6-OHDA-induced dopaminergic neurodegeneration, in contrast to the calnexin CNX-1 and the putative inositol phosphate receptor ITR-1. CRT-1 is a calcium-binding and calcium-storing chaperone of the endoplasmic reticulum (ER) which is predominantly expressed in the *C. elegans* intestine and induced by organismal stress [44]. CRT-1 is required for neuronal cell death induced by hyperactivated MEC-4(d) (<u>mec</u>hanosensory abnormality) and DEG-1(d) (<u>deg</u>eneration of certain neurons) ion channels, as well as by a constitutively active Gas subunit [39]. Furthermore, *crt-1* mutation alleviates dopaminergic cell death caused by gain-of-function mutations of the TRP-4 (<u>t</u>ransient <u>r</u>eceptor <u>p</u>otential) channel [40]. ITR-1 and CNX-1 are less well characterised than CRT-1, but mutation of *itr-1* also reduces neuron loss as induced by hyperactivity of MEC-4(d) and TRP-4, albeit to a lesser extent than mutation of *crt-1* [39,40]. *cnx-1* mutation only suppressed MEC-4(d), but not TRP-4-induce cell death [39,40]. CRT-1, ITR-1 and CNX-1 are thought to be required for calcium release from the ER that is needed for necrotic cell death [39]. When necrosis is linked to increased calcium influx across the plasma membrane, as conferred by the hyperactivated DEG-3(d) channel, necrosis is not blocked by *crt-1* mutation [39]. Finally, CRT-1 also appears to influence the regeneration of axons after laser cut for ALM neurons [45], but not PLM neurons [46]. In the ALM axon regeneration model, the CED-3 and CED-4 apoptosis proteins also promote neuronal repair [45]. Is it possible to reconcile the differential influence of *crt-1* mutation on different types of necrosis and axonal repair with the increased sensitivity to 6-OHDA-induced dopaminergic neurodegeneration? It has been speculated that depending on the levels and duration of the calcium signal, distinct pathways driving neuronal regeneration or cell death can be triggered [47]: Low calcium transients would promote neuronal repair, whereas high levels of increased calcium over a longer period of time would trigger necrosis. 6-OHDA might thus induce a different injury as compared to locally applied laser cuts or the aforementioned necrosis models.

The apoptosis pathway and calreticulin are conserved between *C. elegans* and humans. It is thus conceivable that both apoptosis and necrosis play a role in neurodegeneration in Parkinson’s disease. To the best of our knowledge, apoptosis and necrosis genes have not been defined as Mendelian Parkinson’s loci, and have not been associated with an increased risk of Parkinson’s disease based on genome wide association studies. However, it is noteworthy that the inheritance of Parkinson’s disease is considered to be comparably low, and that the vast majority of disease cases appear to be defined by environmental factors. Thus, apoptosis or oxidative stress signalling linked to apoptosis proteins, as well as necrosis, could well have an important role in protecting dopaminergic neurons in idiopathic Parkinson’s disease.

## Methods

### *C. elegans* strains and maintenance

Strains were grown at 20°C. *wdr-23(tm1817)* and *tsp-17(tm4995)* mutants were generated and kindly provided by Shohei Mitani of the National Bioresource Project (NBRP) (http://shigen.nig.ac.jp/c.elegans/). Additional information about alleles in this study is provided by WormBase (www.wormbase.org). Strains from the *Caenorhabditis* Genetics Center (CGC) or the NBRP were outcrossed to the BY200 strain to eliminate unlinked mutations and to introduce a green fluorescent protein (GFP) marker for dopaminergic neurons. The isolated *gt1981* mutant was outcrossed six times to the BY200 wild-type strain.

**BY200** *vtIs1[pdat-1∷gfp; rol-6*] V
**TG4100** *vtIs1* V; *glit-1(gt1981)* X
**TG4101** *vtIs1* V; *glit-1(gk384527)* X
**TG4102** *vtIs1* V; *glit-1(ok237)* X
**TG2400** *dat-1(ok157)* III; *vtIs1* V
**TG4105** *dat-1(ok157)* III; **vtIs1* V; glit-1(gt1981)* X
**N2 Bristol**
**TG4111** *skn-1(zu67) IV/nT1 [unc-?(n754) let-?*] (IV;V); *vtIs1* V;
**TG4113** *skn-1(lax120); vtIs1* V;
**TG4114** *skn-1(lax120); vtIs1* V; *glit-1(gt1981)* X
**TG4115** *wdr-23(tm1817)* I; *vtIs1* V;
**TG4116** *wdr-23(tm1817)* I; *vtIs1* V; *glit-1(gt1981)* X
**TG4120** *pmk-1(km25)* IV; *vtIs1* V
**TG4122** *jnk-1(gk7)* IV; *vtIs1* V
**TG4124** *pmk-3(ok169)* IV; *vtIs1* V
**TG4126** *kgb-1(ok169)* IV; *vtIs1* V;
**TG4127** *kgb-1(ok169)* IV; *vtIs1* V; *glit-1(gt1981)* X
**TG4128** *pmk-1(km25)* IV; *vtIs1* V; *glit-1(gt1981)* X
**TG4129** *jnk-1(gk7)* IV; *vtIs1* V; *glit-1(gt1981)* X
**TG4130** *pmk-3(ok169)* IV; *vtIs1* V; *glit-1(gt1981)* X
**TG2436** *vtIs1* V; *tsp-17(tm4995)* X
**TG4131** *vtIs1* V; *tsp-17(tm4995)* X; *glit-1(gt1981)* X
**TG4135** *vtIs1* V; *nlg-1(ok259)* X
**TG4136** *vtIs1* V; *glit-1(gt1981)* X; *nlg-1(ok259)* X
**TG4137** *vtIs1* V; *nrx-1(ok1649)* X
**TG4138** *vtIs1* V; *glit-1(gt1981)* X; *nrx-1(ok1649)* X
**TG4139** *vtIs1* V; *nrx-1(wy778)* X
**TG4140** *vtIs1* V; *glit-1(gt1981)* X; *nrx-1(wy778)* X
**TG2410** *vtIs1* V; *dop-1(vs100)* X
**TG4141** *vtIs1* V; *dop-1(vs100)* X; *glit-1(gt1981)* X
**TG2412** *dop-2(vs105)* V; *vtIs1* V
**TG4142** *dop-2(vs105)* V; *vtIs1* V; *glit-1(gt1981)* X
**TG2414** *vtIs1* V; *dop-3(vs106)* X
**TG4143** *vtIs1* V; *glit-1(gt1981)* X; *dop-3(vs106)* X
**TG4144** *dop-2(vs105)* V; *vtIs1* V; *dop-1(vs100)* X
**TG4145** *dop-2(vs105)* V; *vtIs1* V; *dop-1(vs100)* X; *glit-1(gt1981)* X
**TG4146** *vtIs1* V; *dop-1(vs100)* X; *dop-3(vs106)* X
**TG4147** *vtIs1* V; *dop-1(vs100)* X; *glit-1(gt1981) X; dop-3(vs106)* X
**TG2466** *dop-2(vs105)* V; *vtIs1* V; *dop-3(vs106)* X
**TG4148** *dop-2(vs105)* V; *vtIs1* V; *glit-1(gt1981); dop-3(vs106)* X
**TG2415** *dop-2(vs105)* V; *vtIs1* V; *dop-1(vs100)* X; *dop-3(vs106)* X
**TG4149** *dop-2(vs105)* V; *vtIs1* V; *dop-1(vs100)* X; *glit-1(gt1981) X; dop-3(vs106)* X
**TG2396** *bas-1(ad446)* III; *vtIs1* V
**TG4150** *bas-1(ad446)* III; *vtIs1* V; *glit-1(gt1981)* X
**TG2399** *vtIs1* V; *cat-1(ok411)* X
**TG4151** *vtIs1* V; *glit-1(gt1981)* X; *cat-1(ok411)* X
**TG2395** *cat-2(e1112)* II; *vtIs1* V;
**TG4152** *cat-2(e1112)* II; *vtIs1* V; *glit-1(gt1981)* X
**TG4153** *unc-119(ed3)* III; cxTi10816 IV; *otIs433* [*dat-1∷NLS∷RFP ttx-3∷mCh*] V; Ex[P*glit-1∷g/p*] a
**TG4094** *unc-119(ed3)* III; cxTi10816 IV; *otIs433* [*dat-1∷NLS∷rfp ttx-3∷mCh*] V; Ex[P*glit-1∷g/p*] b
**TG4095** *unc-119(ed3)* III; cxTi10816 IV; *otIs433* [*dat-1∷NLS∷rfp ttx-3∷mCh*] V; Ex[P*glit-1∷g/p*] c
**TG4162** *crt-1(ok948)* V; *vtIs1* V;
**TG4163** *ced-4(n1162)* III; *vtIs1* V;
**TG4165** *ced-4(n1162)* III; *vtIs1* V; *glit-1(gt1981)* X
**TG4166** *ced-3(n717)* IV; *vtIs1* V;
**TG4167** *ced-3(n717)* IV; *vtIs1* V; *glit-1(gt1981)* X
**TG4168** *vtIs1* V; *ced-13(sv32)* X;
**TG4169** *vtIs1* V; *glit-1(gt1981) X; ced-13(sv32)* X;
**TG4171** vtIs1V; CB4856
**TG4096** *ttTi5605* II; *unc-119(ed9)* III; *otIs433* [*dat-1∷NLS∷rfp ttx-3∷mCh*] V; *Ex[Pglit-1∷gfp∷glit-1*] a
TG4097 *ttTi5605* II; *unc-119(ed9)* III; *otIs433* [*dat-1∷NLS∷* rfp *ttx-3∷mCh*] V; *Ex[Pglit-1∷gfp∷glit-1*] b
**BY200** *vtIs1[pdat-1∷gfp; rol-6*] V
**N2 Bristol**
**TG4098** *ttTi5605* II; *unc-119(ed9)* III; *otIs433* [*dat-1∷NLS∷rfp ttx-3∷mCh*] V; *Ex[Pglit-1∷gfp∷glit-1*]
**TG2395** *cat-2(e1112)* II; *vtIs1* V;
**TG2396** *bas-1(ad446)* III; *vtIs1* V
**TG2399** *vtIs1* V; *cat-1(ok411)* X
**TG2400** *dat-1(ok157)* III; *vtIs1* V
**TG2410** *vtIs1* V; *dop-1(vs100)* X
**TG2412** *dop-2(vs105)* V; *vtIs1* V
**TG2414** *vtIs1* V; *dop-3(vs106)* X
**TG2415** *dop-2(vs105)* V; *vtIs1* V; *dop-1(vs100)* X; *dop-3(vs106)* X
**TG2436** *vtIs1* V; *tsp-17(tm4995)* X
**TG2466** *dop-2(vs105)* V; *vtIs1* V; *dop-3(vs106)* X
**TG2477** **vtIs1* V; dop-3(vs106) X; tsp-17(tm4995)* X
**TG4094** *unc-119(ed3)* III; cxTi10816 IV; *otIs433* [*dat-1∷NLS∷RFPrfp ttx-3∷mCherry*] V; Ex[P*glit- 1∷gfp∷3′glit-1 Pmyo-3∷mCherry*] b
**TG4095** *unc-119(ed3)* III; cxTi10816 IV; *otIs433* [*dat-1∷NLS∷RFPrfp ttx-3∷mCherry*] V; Ex[P*glit- 1∷gfp∷3′glit-1 Pmyo-3∷mCherry*] c
**TG4096** *ttTi5605* II; *unc-119(ed9)* III; *otIs433* [*dat-1∷NLS∷rfp ttx-3∷mCh*] V; Ex[P*glit-1∷gfp∷glit- 1∷3′glit-1; Pmyo-2∷mCherry*] a
**TG4097** *ttTi5605* II; *unc-119(ed9)* III; *otIs433* [*dat-1∷NLS∷ rfp ttx-3∷mCh*] V; Ex[P*glit-1∷gfp∷glit- 1∷3′glit-1; Pmyo-2∷mCherry*] b
**TG4098** *ttTi5605* II; *unc-119(ed9)* III; *otIs433* [*dat-1∷NLS∷rfp ttx-3∷mCh*] V; Ex[P*glit-1∷gfp∷glit- 1∷3′glit-1; Pmyo-2∷mCherry*] c
**TG4099** *ttTi5605* II; *Is[Pglit-1∷glit-1∷3′glit-1*, integration into *ttTi5605* MosSCI site] II; *unc-119(ed9)* III; *vtIs1* V; *glit-1(gt1981)* X
**TG4100** *vtIs1* V; *glit-1(gt1981)* X
**TG4101** *vtIs1* V; *glit-1(gk384527)* X
**TG4102** *vtIs1* V; *glit-1(ok237)* X
**TG4105** *dat-1(ok157)* III; **vtIs1* V; glit-1(gt1981)* X
**TG4111** *skn-1(zu67) IV/nT1 [unc-?(n754) let-?*] (IV;V); *vtIs1* V;
**TG4113** *skn-1(lax120); vtIs1* V;
**TG4114** *skn-1(lax120); vtIs1* V; *glit-1(gt1981)* X
**TG4115** *wdr-23(tm1817)* I; *vtIs1* V;
**TG4116** *wdr-23(tm1817)* I; *vtIs1* V; *glit-1(gt1981)* X
**TG4120** *pmk-1(km25)* IV; *vtIs1* V
**TG4122** *jnk-1(gk7)* IV; *vtIs1* V
**TG4124** *pmk-3(ok169)* IV; *vtIs1* V
**TG4126** *kgb-1(ok169)* IV; *vtIs1* V;
**TG4127** *kgb-1(ok169)* IV; *vtIs1* V; *glit-1(gt1981)* X
**TG4128** *pmk-1(km25)* IV; *vtIs1* V; *glit-1(gt1981)* X
**TG4129** *jnk-1(gk7)* IV; *vtIs1* V; *glit-1(gt1981)* X
**TG4130** *pmk-3(ok169)* IV; *vtIs1* V; *glit-1(gt1981)* X
**TG4131** *vtIs1* V; *tsp-17(tm4995)* X; *glit-1(gt1981)* X
**TG4135** *vtIs1* V; *nlg-1(ok259)* X
**TG4136** *vtIs1* V; *glit-1(gt1981)* X; *nlg-1(ok259)* X
**TG4137** *vtIs1* V; *nrx-1(ok1649)* X
**TG4138** *vtIs1* V; *glit-1(gt1981)* X; *nrx-1(ok1649)* X
**TG4139** *vtIs1* V; *nrx-1(wy778)* X
**TG4140** *vtIs1* V; *glit-1(gt1981)* X; *nrx-1(wy778)* X
**TG4141** *vtIs1* V; *dop-1(vs100)* X; *glit-1(gt1981)* X
**TG4142** *dop-2(vs105)* V; *vtIs1* V; *glit-1(gt1981)* X
**TG4143** *vtIs1* V; *glit-1(gt1981)* X; *dop-3(vs106)* X
**TG4144** *dop-2(vs105)* V; *vtIs1* V; *dop-1(vs100)* X
**TG4145** *dop-2(vs105)* V; *vtIs1* V; *dop-1(vs100)* X; *glit-1(gt1981)* X
**TG4146** *vtIs1* V; *dop-1(vs100)* X; *dop-3(vs106)* X
**TG4147** *vtIs1* V; *dop-1(vs100)* X; *glit-1(gt1981) X; dop-3(vs106)* X
**TG4148** *dop-2(vs105)* V; *vtIs1* V; *glit-1(gt1981); dop-3(vs106)* X
**TG4149** *dop-2(vs105)* V; *vtIs1* V; *dop-1(vs100)* X; *glit-1(gt1981) X; dop-3(vs106)* X
**TG4150** *bas-1(ad446)* III; *vtIs1* V; *glit-1(gt1981)* X
**TG4151** *vtIs1* V; *glit-1(gt1981)* X; *cat-1(ok411)* X
**TG4152** *cat-2(e1112)* II; *vtIs1* V; *glit-1(gt1981)* X
**TG4153** *unc-119(ed3)* III; cxTi10816 IV; otIs433 [*dat-1∷NLS∷RFP ttx-3∷mCherry*] V; Ex[Pglit- *1∷gfp∷3′glit-1; Pmyo-3∷mCherry*] a
**TG4162** *crt-1(ok948)* V; *vtIs1* V;
**TG4163** *ced-4(n1162)* III; *vtIs1* V;
**TG4165** *ced-4(n1162)* III; *vtIs1* V; *glit-1(gt1981)* X TG4166 *ced-3(n717)* IV; *vtIs1* V;
**TG4167** *ced-3(n717)* IV; *vtIs1* V; *glit-1(gt1981)* X
**TG4168** *vtIs1* V; *ced-13(sv32)* X;
**TG4169** *vtIs1* V; *glit-1(gt1981) X; ced-13(sv32)* X;
**TG4171** vt I s 1 V; CB4856
**TG4176** *ttTi5605* II; *Is[Pdat-1∷glit-1∷3′glit-1*, integration into *ttTi5605* MosSCI site] II; *unc-119(ed9)* III; *vtIs1* V; *glit-1(gt1981)* X
**TG4177** *ttTi5605* II; *Is[Ppha-4∷glit-1∷3′glit-1*, integration into *ttTi5605* MosSCI site] II; *unc-119(ed9)* III; *vtIs1* V; *glit-1(gt1981)* X
**TG4178** *ttTi5605* II; *Is[Pelt-2∷glit-1∷3′glit-1*, integration into *ttTi5605* MosSCI site] II; *unc-119(ed9)* III; *vtIs1* V; *glit-1(gt1981)* X
**TG4344** *ttTi5605* II; *unc-119(ed9)* III; *vtIs1* V; *Ex[Pglit-1∷glit-1∷3′glit-1, unc-119(+), Pmyo-2∷mCherry, Pmyo-3∷mCherry*] a
**TG4345** *ttTi5605* II; *unc-119(ed9)* III; *vtIs1* V; *Ex[Pglit-1∷glit-1∷3′glit-1, unc-119(+), Pmyo-2∷mCherry, Pmyo-3∷mCherry*] b
**TG4346** *egl-1(n1084n3082) V; vtIs1* V
**TG4347** *ced-9(n1950)* III; *vtIs1* V
**TG4348** *daf-2(e1368)* III; *vtIs1* V
**TG4349** *daf-2(e1368)* III; *vtIs1* V; *glit-1(gt1981)* X
**TG4350** *daf-16(mu86)* I; *vtIs1* V
**TG4351** *daf-16(mu86)* I; *vtIs1* V; *glit-1(gt1981)* X
**TG4352** *cnx-1(nr2009)* III; *vtIs1* V
**TG4353** *itr-1(sa73)* IV; *vtIs1* V

### Mutagenesis and mapping

L4 and young adult staged BY200 animals were mutagenised with 25 mM ethyl methanesulfonate (EMS) in M9 buffer for 4 hours at 20°C, washed and grown at 15°C overnight on NGM plates seeded with OP50 *E. coli*. The animals were bleached once they started to lay eggs and the F1 progeny let hatch overnight in M9. This synchronised F1 population was left to develop on seeded plates until reaching early adult stage. The F1 adults were then washed off in M9 again to lay a synchronised population of F2 L1 larvae which was treated with 10 mM 6-OHDA and screened for dopaminergic neuron loss after 72 h. Candidates were transferred to separate plates, backcrossed at least three times and re-tested for their hypersensitivity to 6-OHDA. The genomic DNA was sent for whole- genome sequencing and SNP mapping was performed as previously described [14,15].

### Generation of transgenic animals and imaging

For generation of transgenic constructs, genomic DNA was amplified using primers containing additional 8 base pair restriction sites (marked in italics) for insertion into the pCFJ151 vector [48]. In the following primer sequences, restriction sites are highlighted in italics, translation start and translation stop codons in bold and codons are separated by a space. Primers for the transcriptional reporter construct *AscI_Pglit-1_NotI_* GFP_*FseI*_3′glit-1_*PacI*: *AscI*_Pglit-1 – GCTA*ggcgcgcc*GTATCTGGCATTGGCTCG; Pglit-1_*NotI* – GCTA *gcg gcc gca* **CAT**TCCATGTGACGCGAT; *NotI*_GFP – GCTAgc *ggc cgc* AGT AAA GGA GAA GAA CTT TTC ACT GG; *GFP_FseI-* AAGTTA *ggc cgg ccc* CTT GTA TGG CCG GCT AG; *FseI_3′glit-1-* AC AAG *ggg ccg gcc* **TAA**CTTTCAAAGTTTGTAAATAATGTATAATTTA; 3′glit-1_*Pac* – GCTAttaattaaCCAGTTGCAGTGTTTTTTTG. Further primers for the GFP-tagged translational constructs *AscI*_Pglit-1_*NotI*_GFP_*FseI*_glit-1_3′glit-1_*PacI*: *FseI*_glit-1_3′*glit-1* – GG CCA TAC AAG G*gg ccg gcc* ATG TTC ACC GGC ACA ATT; glit-1_3′glit-1_*PacI*-TAGGGCCCTCAA*ttaattaa*GACAATACATGTTTTCTTTTTGAAAATG. Additional primers for the untagged translational constructs *AscI*_Pglit-1_*FseI*_glit-1_3′*glit-1*_*PacI* and *AscI*_Pdat-1_*FseI*_glit-1_3′*glit- 1*_*PacI*:*AscI*_Pglit-1 – AGATTA*ggcgcgcc*GTATCTGGCATTGGCTCG; Pglit-1_*FseI* – AGAAAAggc *cgg cc***a CAT**T CCAT GT GACGCGAT; *AscI*_Pdat-1 – ATT A*ggcgcgcc* AATGTTT CTAGTCGTTTTTGTATTTTAAAG; Pdat-1_*FseI*- AGAAAA*ggc cgg cc*a CATGGCTAAAAATTGTTGAGATTCG. The *glit-1* coding sequence was amplified from cDNA (the AT-rich *glit-1* introns rendered cloning unfeasible). The vector was adapted to include the following restriction site architecture: *AscI*_SgfI_*NotI*_FseI_*PacI*. The strains were generated by microinjections of *unc-119(ed3)* mutants. For microscopy, animals were transferred into a drop of 25 mM levamisole solution on top of a 2.5% (w/v) agar pad on a microscope slide. Images were acquired using a DeltaVision microscope (Applied Precision) and analysed with the softWoRx Suite (Applied Precision) and ImageJ [49].

### *C. elegans* assays

All *C. elegans* assays were performed in a blinded manner.

To obtain synchronized L1 larvae for oxidative stress assays, 1-10 adult animals were incubated to lay eggs in 70 μl M9 without food on a benchtop shaker at 20°C, 500 rpm for 24-30 h. Due to the swimming-defect of *tsp-17(tm4995)*, all L1 larvae for experiments that included this strain were obtained by filtering a mixed population of animals. For **6-OHDA assays**, 10 μL 200 mM ascorbic acid and 10 μL of the respective 6-OHDA 5 x stock concentration were added to 30 μL of L1-stage larvae in M9. 6-OHDA concentration was chosen such that differential sensitives of various strains could be efficiently analysed. After 1 hour incubation at 20°C and shaking at 500 rpm, 150 μL M9 buffer was added to oxidise and inactivate the 6-OHDA. The animals were pipetted to one half of an NGM plate containing a stripe of OP50 bacteria on the opposite half of the plate and adult animals and eggs were removed to prevent growth of animals that were not treated at L1 stage. Plates were incubated at 20°C before blind scoring of the 6-OHDA- induced degeneration using a Leica fluorescent dissecting microscope. Unless indicated otherwise, animals were scored 72h after treatment in technical duplicates. For treatment of different developmental stages, the respective stages were selected from a mixed population of animals after 6-OHDA treatment. Replicates for strains shown in the same graph were performed in parallel.

For **paraquat and H_2_O_2_ assays**, 10 μL of the respective 5x stock concentration was added to 40 μL of L1-stage larvae in M9. After 1 hour incubation in the shaker 150 μL M9 buffer was added and the L1 larvae were counted after pipetting them to a seeded NGM plate. Surviving animals were scored again after 24 h.

**Dye leakage assays** were performed as described previously [19,20]. A mixed, well-fed population of animals was washed off the plate and incubated for 3 hours in 2.5% (wt/vol) Brilliant Blue FCF (Sigma) and 2.5 % (wt/vol) fluorescein sodium salt (Sigma), respectively. The dyes were solubilised in a saturated solution of OP50 *E. coli* in LB (<u>L</u>ysogeny <u>B</u>roth) medium. After incubation, the animals were pipetted onto NGM plate seeded with OP50 *E. coli*. For microscopy, single animals were transferred immediately into a drop of 25 mM levamisole solution on top of a 2.5% (w/v) agar pad on a glass slide. For the blue dye (‘Smurf’) assay, images were taken using a camera mounted to a Leica stereo microscope. To determine fluorescein uptake, a bright-field image and a fluorescence image using the GFP channel were taken with a Zeiss Axio Scope. Bright-field and fluorescence image channels were merged using ImageJ [49].

For lifespan assays, 50 L4 stage animals were selected per strain and transferred to new plates using a platinum wire every day during the first 6 days and then only to avoid mixing with the progeny. Animals that did not move after touch with a platinum pick were scored as dead. Dead animals were counted and removed every day. Bag of worms, dry animals and burst animals were censored. The data was analysed using OASIS (Online application for Survival Analysis, http://sbi.postech.ac.kr/oasis/) [50].

**Dopamine paralysis assays** were carried out as previously described [51]. Staged adult animals were incubated for 20 minutes on a plate containing dopamine and assessed for their ability to complete body movement through minimum and maximum amplitude during a 5 second observation period. Two plates containing 25 animals each were assessed for each condition.

For the **swimming-induced paralysis (SWIP)** assay 5-15 L4 stage larvae were transferred into a glass well in 40 μL water. The number of paralysed animals scored using a Leica dissecting microscope every minute for 30 minutes [25].

**Basal slowing assays** were performed as previously described [52]. NGM plates were prepared and half of them seeded with HB101 bacteria before incubating them at 37°C overnight. Also, mid-L4 larvae were transferred on separate plates. The next day, these staged young adult animals were put in 40 μL M9 for 2 minutes to clean them from bacteria and then transferred to the centre of a plate. 6 animals each were put on a plate with and without bacteria, respectively. After a 2 minutes adaptation time, the locomotion rates of each animal was quantified by counting the number of body bends completed in five consecutive 20 s intervals. The experiment was performed twice on different days.

### qRT-PCR

L1 stage animals were filtered from two 9 cm plates containing mixed stage animals. Animals were washed twice and then split in half for control treatment (addition of ascorbic acid and water only) and 6-OHDA treatment. After treatment, RNA extraction was performed based on He, F. (2011). Total RNA Extraction from *C. elegans*. Bio-protocol Bio101: e47. DOI: 10.21769/BioProtoc.47. The following changes were introduced: The chloroform extraction was repeated once more and 0.1 volumes of 3 M NaAc were added to the propanol extraction to help RNA precipitation. The RNA pellet was resuspended in 15μL H2O. The cDNA was digested using the Turbo DNA-free kit (Ambion Life Technologies) in 18 μL total volume. 1 μg of RNA was reverse transcribed to cDNA with 15 bp oligo dT primers using the M-MLV Reverse Transcriptase (Promega) according to manufacturers’ instructions. The cDNA was analysed by qRT-PCR using the StepOnePlus Real-Time PCR System (Applied Biosystems) applying melting temperature of 62°C. The following primers were designed using QuantPrime [53]: K08F4.7_F: AT GGT CAAAGCT GAAGCCAACG and K08F4.7_R: ACTGACCGAATTGTTCTCCATCG. The PCR mixture consisted of 0.2 μM primers, cDNA (diluted 1:50) and 1x Power SYBR Green PCR Master Mix (Thermo Fisher Scientific). All reactions were run in triplicates. Relative abundance was calculated using the ΔΔCt method [54]. To control for template levels, values were normalised against the control genes *Y45F10D.4* and *pmp-3* [55,56].

### Bioinformatics analysis

**GLIT-1 extra- and intracellular and transmembrane domains and signal peptides** were predicted with Phobius (http://phobius.sbc.su.se/) [57].

**GLIT-1 protein structure model** was calculated in SWISS-MODEL (<u>swissmodel.expasy.org)</u> [58] and visualised with Swiss-PdbViewer 4.1.0 (http://www.expasy.org/spdbv/) [59].

For **phylogenetic analysis of GLIT-1**, NCBI Protein BLAST (Basic Local Alignment Search Tool) was used to search the ‘reference proteins’ database of the indicated organism against the GLIT-1 protein sequence. A multiple alignment was performed using COBALT with the 250 best-matching sequences from all organisms.

Protein alignments to **visualise conservation** of GLIT-1 domains and residues alignment of protein sequences was performed in CLC workbench Version 7.6.4 using the default parameters: Gap open cost 10.0, Gap extension cost 1.0, End gap cost as any other, Alignment very accurate (slow) and create a the tree (Tree construction method: Neighbour Joining, Protein distance measure: Jukes- Cantor, Bootstrapping: 100 Replicates).

### Statistical analysis

The **two-tailed t-test** and were performed with the Analysis ToolPak Add-In in Microsoft Excel 2010. To choose a t-test assuming equal or unequal variances, respectively, the same Add-in was used to conduct an F-test.

To compare 6-OHDA treatment data, a **G-Test** was performed with R Studio 1.0.44 DescTools descriptive statistics tools package. p-values were calculated for each biological replicate and for the pooled biological replicate data, and the least significant of these p-values is indicated in the graphs. Unless otherwise indicated, data is only marked as significant if all replicates and the pooled replicate data were found to be significant.

## Acknowledgements

We thank Neda Masoudi for advice at initial stages of the project, Radoslav Lukoszek for advising on qRT-PCR experiments, Bjoern Schumacher for sending *egl-1* and *ced-9* strains, Ulrike Gartner for reading the manuscript and WormBase. We would like to thank Oliver Hobert for sharing the *otIs433* marker strain. Some strains were provided by the CGC, which is funded by NIH Office of Research Infrastructure Programs (P40 0D010440).

## Supporting information

**F1 Fig. *gt1981* mutant characterisation.**

(A) Developmental stages of wild-type and *gt1981* mutant L1 stage larvae 48 hours after treatment with and without 10 mM 6-OHDA. *C. elegans* develops via the L1, L2 (in red), L3 (in orange), L4 (green) and young adult stage (in light blue) into adults (in dark blue). Error bars = SEM of 3 biological replicates, each with 60-110 animals per treatment and strain. Total number of animals per condition n = 215-270 (n.s. p>0.05; G-Test). (B) Dopaminergic head neurons 72 hours after treatment with 10 mM 6-OHDA in BY200 wild-type animals, *glit-1(gt1981)* and *glit-1(gk384527)* homozygous mutants and *glit-1/glit-1(gk384527)* transheterozygote mutant animals. Error bars = SEM of 2-3 experiments, each with 60-200 animals per strain. Total number of animals per strain n = 270-340 (n.s. p>0.05; G-Test). (C) Dopaminergic head neurons 72 hours after treatment with 10 mM 6-OHDA in BY200 wild-type animals and *glit-1(gt1981)* homozygous and *glit-1(gt1981)/+* heterozygous mutants. Error bars = SEM of 2 experiments, each with 30-105 animals per strain. Total number of animals per strain n = 65-210 (****p<0.0001, n.s. p>0.05; G-Test). (D) Effect of *dat-1* mutation on dopaminergic neurodegeneration after treatment with 10 mM 6-OHDA. Error bars = SEM of 2 experiments, each with 30-110 animals per strain. Total number of animals per strain n = 100-400 (****p<0.0001; G-Test).

**S2 Fig. GLIT-1 protein architecture.**

(A) Predicted probabilities for the occurrence of a signal peptide (in red), a transmembrane domain (in grey), a cytoplasmic part (in green) and a non-cytoplasmic part (in blue) as calculated with Phobius (http://phobius.sbc.su.se/).

**S3 Fig. GLIT-1 phylogenetic alignment.**

(A) Phylogenetic tree of GLIT-1 aligned to protein blast matches of selected species (listed below). The root of the tree is marked in green, esterases are marked in blue and vertebrate and invertebrate neuroligin groups are framed. The subtree containing GLIT-1 is indicated with a red asterisk. (B) Blow-up of the GLIT-1-containing subtree. Acyrthosiphon pisum (pea aphid), *Caenorhabditis (C.) briggsae* (nematode), *Ciona intestinalis* (sea squirt), *Culex quinquefasciatus* (southern house mosquito), *Danio rerio* (zebrafish), *Drosophila melanogaster* (fruit fly), *Homo sapiens* (human), *Ixodes scapularis* (tick), *Mus musculus* (house mouse), *Nasonia vitripennis* (parasitoid wasp), *Pediculus humanus corporis* (body louse), *Schistosoma mansoni* (parasitic trematode), *Strongylocentrotus purpuratus* (purple sea urchin), *Tribolium castaneum* (red flour beetle), *Trichinella spiralis* (parasitic nematode).

**S4 Fig. GLIT-1 domain conservation.**

Alignment of acetylcholinesterases (ace) and neuroligins (nlg) from mouse *(Mus musculus*, Mm), human *(Homo sapiens*, Hs), fruit fly *(Drosophila melanogaster*, Dm), and *C. elegans* (Ce). Carboxyesterase-like domain and transmembrane domain are indicated with blue arrows on top of the GLIT-1 sequence. The conservation score is indicated with a bar graph at the bottom of the alignment.

**S5 Fig. GLIT-1 amino acid conservation.**

Magnification of alignment of acetylcholinesterases (ace) and neuroligins (nlg) from mouse (*Mus musculus*, Mm), human *(Homo sapiens*, Hs), fruit fly *(Drosophila melanogaster*, Dm), and *C. elegans* (Ce) to show the (A) proline residue mutated in *glit-1(gt1981)* (P113L) and the (B) serine, (C) histidine and (D) glutamate residues that form part of the catalytic triad of acetylcholinesterases. High conservation is highlighted in red and low conservation in blue. The conservation score is indicated with a bar graph at the bottom of the alignment.

**S6 Fig. *glit-1* expression.**

(A) L3 stage larva expressing *Ex[Pglit-1∷gfp∷glit-1*]. (B) Embryo and L4 stage larva expressing *Ex[Pglit-1∷gfp∷glit-1*]. (C) – (E) Head region of the L4 stage larva from Fig 2B-F. The green channel shows expression of *Ex[Pglit-1∷gfp∷glit-1*]. The red channel shows expression of Is[Pdat- *1∷NLS∷rfp;Pttx-3∷mCherry*] for labelling of dopaminergic neuron nuclei, as well as the pharynx muscle marker Ex[*Pmyo-2∷mCherry*] and the body muscle marker Ex[*Pmyo-3∷mCherry*] that were used for injections. (F) L1 stage larva expressing *Ex[Pglit-1∷gfp*]. (G) and (H) Head region of adult animals. The green channel shows expression of *Ex[Pglit-1∷gfp*]. The red channel shows expression of *Is[Pdat-1∷NLS∷rfp;Pttx-3∷mCherry*] for labelling of dopaminergic neuron nuclei, as well as the pharynx muscle marker *Ex[Pmyo-2∷mCherry*] and the body muscle marker *Ex[Pmyo-3∷mCherry*] that were used for injections.

**S7 Fig. 6-OHDA-induced dopaminergic neurodegeneration in animals carrying *glit-1* extrachromosomal arrays.**

(A) Effects of *Ex[Pglit-1∷gfp∷glit-1*] expression on dopaminergic neurodegeneration in the *glit-1* mutant after treatment with 10 mM 6-OHDA. Error bars = SEM of 2 biological replicates, each with 20-100 animals per strain and concentration. Total number of animals per strain n = 70-200 (n.s. p>0.05; G-Test). (B) Effect of multi-copy *Ex (Pglit-1∷glit-1*] construct on dopaminergic neurodegeneration after treatment with 25 and 50 mM 6-OHDA. Error bars = SEM of 2 biological replicates, each with 80-110 animals per strain. Total number of animals per condition n = 185-215 (n.s. p>0.05,; G-Test). (C) *tsp-17* and (D) *glit-1* mRNA levels in wild-type and mutant L1 stage larvae after 1 hour treatment with 10 mM 6-OHDA. The data are normalised to the control gene *Y45F10D.4*. The average and the respective values for 5 biological replicates (biorep a-e) are indicated. Error bars = SEM of 5 biological replicates. (E) *tsp-17* and (F) *glit-1* mRNA levels in wild-type and mutant L1 stage larvae after 1 hour treatment with 10 mM 6-OHDA. The data are normalised to the control gene *pmp-3*. The average and the respective values for 5 biological replicates (biorep a-e) are indicated. Error bars = SEM of 5 biological replicates.

**S8 Fig. Uptake of fluorescein sodium salt in wild-type and *glit-1* mutant animals.**

Overlay of fluorescence image (in green) and bright-field image (in grey) of wild-type and *glit- 1(gt1981)* mutant animals after 3 hours of incubation in fluorescein sodium salt.

**S9 Fig. Uptake of Brilliant Blue FCF (’Smurf’ assay) in wild-type and *glit-1* mutant animals.**

Wild-type and *glit-1* mutant animals after 3 hours of incubation in Brilliant Blue FCF.

**S10 Fig. Effect of various 6-OHDA concentrations and neuroligin and neurexin mutations on 6- OHDA-induced dopaminergic neurodegeneration.**

(A) Dopaminergic head neurons 72 hours after treatment with 10 mM 6-OHDA in wild-type and *glit-*1 mutant animals. Error bars = SEM of 1-2 experiments, each with 90-115 animals per strain. Total number of animals per strain n = 115-220. (B) Effects of *nlg-1* and *nrx-1* mutations on dopaminergic neurodegeneration after treatment with 0.75 mM, (C) 10 mM and (D) 50 mM 6-OHDA. Error bars = SEM of 2 biological replicates, each with 50-110 animals per strain and concentration. Total number of animals per condition n = 150-220 (****p<0.0001, **p<0.01, *p<0.05, n.s. p>0.05; G-Test).

**S11 Fig. *glit-1* genetically interacts with dopamine metabolism enzymes.**

(A) Effect of dopamine receptor mutations on dopaminergic neurodegeneration after treatment with 0.75 mM 6-OHDA. Error bars = SEM of 3 biological replicates, each with 60-115 animals per strain and concentration. Total number of animals per condition n = 180-340 (****p<0.0001, ***p<0.001, **p<0.01, *p<0.05, n.s. p>0.05; G-Test comparing glit-1 mutant sensitivity to sensitivity of double and triple mutants). A significant p-value is only indicated if all or all but one replicate were found to be significant. (B) Effect of mutations in dopamine metabolism genes on dopaminergic neurodegeneration after treatment with 0.75 mM and (C) 10 mM 6-OHDA. Error bars = SEM of 3 biological replicates, each with 60-115 animals per strain and concentration. Total number of animals per condition n = 180-325 (****p<0.0001, ***p<0.001, n.s. p>0.05; G-Test comparing *glit-1* to double mutants). (D) Effect of mutations in the dopamine receptor *dop-2* and the dopamine metabolism genes *cat-1* and *cat-2 (abnormal <u>cat</u>echolamine distribution)* on dopaminergic neurodegeneration after treatment with 25 mM and (E) 50 mM 6-OHDA. Error bars = SEM of 3-4 biological replicates, each with 50-115 animals per strain and concentration. Total number of animals per condition n = 240-430 (****p<0.0001,***p<0.001, n.s. p>0.05; G-Test comparing BY200 wild-type to mutant animal data). A significant p-value indicated if all or all but one replicate were found to be significant. (F) Cartoon illustrating dopamine signalling cartoon (adapted from [23]). Dopamine synthesis in dopaminergic (DAergic) neurons starts with tyrosine (Tyr), which is converted to L-DOPA (L-3,4-<u>d</u>ihydr<u>o</u>xyphenyl<u>a</u>lanine) by the tyrosine hydroxylase CAT-2. L-DOPA is then converted to dopamine (DA) by the aromatic amino acid decarboxylase BAS-1 (<u>b</u>iotenic <u>a</u>mine <u>s</u>ynthesis related). Dopamine is packed into vesicles by the vesicular monoamine transporter CAT-1. Postsynaptic dopamine signalling is stimulated by the D1-like <u>dop</u>amine receptor DOP-1 and inhibited by the D2-like <u>dop</u>amine receptor DOP-3. The D2-like receptor DOP-2 can be postsynaptic, or presynaptic and act as an autoreceptor. The D1-like (stimulatory) and the D2-like (inhibitory) dopamine receptors are indicated in blue and red, respectively. Dopamine is transported back into the dopaminergic neurons via the dopamine transporter DAT-1. Cholinergic neurons signal via the neurotransmitter ACh (<u>a</u>cetyl<u>ch</u>oline), which in turn stimulates muscle action.

**S12 Fig. Dopamine-mediated behaviours in *glit-1* and *tsp-17* mutants.**

(A) Animal speed (measured in body bends/20s) on plates without and with food (’+Food’). Indicated is the average speed per young adult animal (grey symbols) and the average speed per strain (black bar). *cat-2 (abnormal <u>cat</u>echolamine distribution)* is a tyrosine hydroxylase mutant and defective in dopamine synthesis. Error bars = SEM of 2 biological replicates, each with 6 animals per strain and state. Total number of animals per condition n = 12 (****p<0.0001, ***p<0.0001, **p<0.0001; twotailed t-test). (B) Percentage of swimming L4 stage animals. Error bars = SEM of 3-4 biological replicates with 12-18 animals per strain. Total number of animals per strain n = 50-60 (*p<0.05, n.s. p>0.05; two-tailed t-test comparing wild-type and mutant data at 30 minute time point).

**S13 Fig. *glit-1* and *tsp-17* mutant lifespan.**

Lifespan data for second biological replicate including 88-109 animals per strain. The inset shows the mean lifespan with the error bars depicting the standard error (****p<0.0001, ***p<0.001; Bonferroni-corrected; Log-Rank Test).

**S14 Fig. mRNA levels of oxidative stress reporters under basal conditions, normalised to the control gene *Y45F10D.4*.**

(A) *gst-4*, (B) *gcs-1* and (C) *gst-1* mRNA levels in mutant L1 stage larvae under control conditions (treatment with H_2_O instead of 6-OHDA). (A)-(C) The fold change is calculated based on mRNA levels in wild-type L1 stage larvae under control conditions. The data are normalised to the control gene *Y45F10D.4* and the average and the respective values for 5-6 biological replicates (biorep a-f) are indicated. Error bars = SEM of 5-6 biological replicates.

**S15 Fig. mRNA levels of oxidative stress reporters normalised to the control gene *pmp-3*.**

(A) *gst-4*, (B) *gcs-1* and (C) *gst-1* mRNA levels in wild-type and mutant L1 stage larvae after 1 hour treatment with 10 mM 6-OHDA. (D) *gst-4*, (E) *gcs-1* and (F) *gst-1* mRNA levels in mutants L1 stage larvae under control conditions (treatment with H_2_O instead of 6-OHDA). The fold change is calculated based on mRNA levels in wild-type L1 stage larvae under control conditions. (A)-(F) The data are normalised to the control gene *pmp-3* and the average and the respective values for 5-6 biological replicates (biorep a-f) are indicated. Error bars = SEM of 5-6 biological replicates (****p<0.0001, **p<0.01, *p<0.05; two-tailed t-test)‥

**S16 Fig. mRNA levels after paraquat exposure**

(A) *tsp-17, glit-1, gst-4, gcs-1, gst-1* and *crt-1* mRNA levels in wild-type and mutant L1 stage larvae after 1 hour treatment with 3 mM 6-paraquat. The data are normalised to the control gene *Y45F10D.4*. (B) *tsp-17, glit-1, gst-4, gcs-1, gst-1* and *crt-1* mRNA levels in wild-type and mutant L1 stage larvae after 1 hour treatment with 3 mM 6-paraquat. The data are normalised to the control gene *pmp-3*. (A) and (B) The average and the respective values for 3-5 biological replicates (biorep a-e) are indicated. Error bars = SEM of 3-5 biological replicates.

**S17 Fig. Effect of *daf-2* and *daf-16* mutations on 6-OHDA-induced dopaminergic neurodegeneration in wild-type and *glit-1* mutant animals.**

(A) Effect of *daf-2* and *daf-16* mutation on dopaminergic neurodegeneration after treatment with 25 mM and (B) 50 mM 6-OHDA. Error bars = SEM of 3 biological replicates with 85-135 scored animals per strain. Total number of animals per strain n = 320-360 (*p<0.05, n.s. p>0.05; G Test). (C) Effect of *daf-2* and *daf-16* mutation on dopaminergic neurodegeneration in wild-type and *glit-1* single and double mutant animals after treatment with 0.75 mM and (D) 10 mM 6-OHDA. Error bars = SEM of 2- 3 biological replicates with 80-130 scored animals per condition. Total number of animals per strain n = 200-340 (n.s. p>0.05; G Test).

**S18 Fig. Effect of mutations in p38 and JNK stress response pathways on 6-OHDA-induced dopaminergic neurodegeneration in wild-type and *glit-1* mutant animals.**

(A) Effect of p38 and JNK stress response pathway mutations on dopaminergic neurodegeneration after treatment with 0.75 mM 6-OHDA. Error bars = SEM of 3 biological replicates, each with 100-120 animals per strain. Total number of animals per strain n = 300-330 (n.s. p>0.05; G-Test). (B) Effect of *pmk-3* mutation on dopaminergic neurodegeneration after treatment with 0.75 mM 6- OHDA. Error bars = SEM of 2-3 biological replicates, each with 50-110 animals per strain. Total number of animals per strain n = 160-300 (n.s. p>0.05; G-Test). (C) Effect of p38 and JNK stress response pathway mutations on dopaminergic neurodegeneration after treatment with 10 mM 6- OHDA. Error bars = SEM of 2-3 biological replicates, each with 40-105 animals per strain. Total number of animals per strain n = 195-315 (n.s. p>0.05; G-Test). (D) Effect of *pmk-3* mutation on dopaminergic neurodegeneration after treatment with 10 mM 6-OHDA. Error bars = SEM of 2-3 biological replicates, each with 30-100 animals per strain. Total number of animals per strain n = 140-270 (n.s. p>0.05; G-Test). (E) Effect of p38 and JNK stress response pathway mutations on dopaminergic neurodegeneration after treatment with 25mM 6-OHDA. Error bars = SEM of 3 biological replicates, each with 100-110 animals per strain. Total number of animals per strain n = 310-330 (n.s. p>0.05; G-Test). (F) Effect of *pmk-1* mutation on dopaminergic neurodegeneration after treatment with 25 mM 6-OHDA. Error bars = SEM of 3 biological replicates, each with 100-135 animals per strain. Total number of animals per strain n = 350-360 (n.s. p>0.05; G-Test).

**S19 Fig. *glit-1, tsp-17, hsp-4* and *hsp-6* mRNA levels after 6-OHDA exposure**

(A) *hsp-4* and (B) *hsp-6* mRNA levels in wild-type and mutant L1 stage larvae after 1 hour treatment with 10 mM 6-OHDA. The data are normalised to the control gene *Y45F10D.4*. The average and the respective values for 5 biological replicates (biorep a-e) are indicated. Error bars = SEM of 5 biological replicates. (C) *hsp-4* and (D) *hsp-6* mRNA levels in wild-type and mutant L1 stage larvae after 1 hour treatment with 10 mM 6-OHDA. The data are normalised to the control gene *pmp-3*. The average and the respective values for 5 biological replicates (biorep a-e) are indicated. Error bars = SEM of 5 biological replicates.

**S20 Fig. Effect of apoptosis pathway mutations on 6-OHDA-induced dopaminergic neurodegeneration in *glit-1* mutant animals.**

(A) Effect of *ced-4* mutation on dopaminergic neurodegeneration after treatment with 0.75 mM 6- OHDA. Error bars = SEM of 5 biological replicates, each with 60-115 animals per strain and concentration. Total number of animals per condition n = 370-530 (n.s. p>0.05; G-Test). (B) Effect of *ced-3* mutation on dopaminergic neurodegeneration after treatment with 0.75 mM 6-OHDA. Error bars = SEM of 3 biological replicates, each with 50-120 animals per strain and concentration. Total number of animals per condition n = 240-320 (n.s. p>0.05; G-Test). (C) Effect of *ced-13* mutation on dopaminergic neurodegeneration after treatment with 0.75 mM 6-OHDA. Error bars = SEM of 3 biological replicates, each with 90-120 animals per strain and concentration. Total number of animals per condition n = 300-330 (n.s. p>0.05; G-Test).

**S21 Fig. Effect of *cnx-1* and *itr-1* mutation on 6-OHDA-induced dopaminergic neurodegeneration and *crt-1* mRNA levels normalised to the housekeeping gene *pmp-3*.**

(A) Effect of *cnx-1* and *itr-1* mutations on dopaminergic neurodegeneration after treatment with 10, 25 and 50 mM 6-OHDA. Error bars = SEM of 3 biological replicates, each with 70-120 animals per strain and concentration. Total number of animals per condition n = 285-325 (n.s. p>0.05; G-Test). (B) *crt-1* mRNA level analysis after 1h of 6-OHDA treatment in wild-type and mutant L1 stage larvae. (C) *crt-1* mRNA level analysis in mutant L1 stage larvae as compared to wild-type animals under control conditions (treatment with H_2_O instead of 6-OHDA). (B) and (C) The data are normalised to the control gene *pmp-3*. The average and the respective values for 5 biological replicates (biorep a-e) are indicated. Error bars = SEM of 5 biological replicates (n.s. p>0.05; two-tailed t-test).

**S1 Movie.** *Pglit-1∷gfp∷3′glit-1* expression (green channel) in adult animal. The red channel shows expression of *\s[Pdat-1∷NLS∷rfp;Pttx-3∷mCherry*] for labelling of dopaminergic neuron nuclei, as well as the pharynx muscle marker *Ex[Pmyo-2∷mCherry*] and the body muscle marker Ex[Pmyo- *3∷mCherry*] that were used for injections. Dopaminergic CEPD, ADE and CEPV neurons can be seen after 2, 5 and 8 seconds, respectively.

**S1 Text. Genetic interactions between *glit-1* and dopamine metabolism**

(A) In *C. elegans*, dopamine can bind to D_1_-type (stimulatory) dopamine receptor DOP-1 and the D_2_-type (inhibitory) dopamine receptors DOP-2 and DOP-3 after release into the synaptic cleft (simplified cartoon in S11F Fig). We found that mutation of *dop-1* and *dop-3* did not alter 6-OHDA sensitivity, while mutation of *dop-2* led to a slight reduction of 6-OHDA-induced dopaminergic neurodegeneration in *glit-1* mutants (S11A Fig). Furthermore, *dop-2* mutation also decreased dopaminergic neuron loss in wild-type animals (S11 D, E Fig) [13] and *tsp-17* mutants [13]. In summary, the rescue of dopamine neuron loss by *dop-2* mutation is only partial and occurs in all tested genetic backgrounds, indicating that the function of *glit-1* is largely unrelated to *dop-2*. (B) Dopamine is synthesised de novo by CAT-2 (abnormal <u>cat</u>echolamine distribution), a dopamine synthesis-specific tyrosine hydroxylase, and the BAS-1 aromatic amino acid decarboxylase, and packed into vesicles by the CAT-1 vesicular monoamine transporter (S11F Fig) [23]. We found that while *glit-1* mutant sensitivity was not modulated by *bas-1* mutation, dopaminergic neurodegeneration of *glit-1* mutants was increased by *cat-1* and *cat-2* mutations (S11B, C Fig). In contrast, in a wild-type background *cat-1* mutation decreased dopaminergic neuron loss and *cat-2* mutation did not show an effect (S11D, E Fig). Mutation of CAT-2, the key dopamine synthesis enzyme, is expected to cause reduced dopamine production. As dopamine and 6-OHDA likely compete for uptake via the dopamine transporter, it is conceivable that overall decreased dopamine levels cause increased neuronal uptake of 6-OHDA, and vice versa. Thus, a *cat-2* mutation might lead to increased 6-OHDA uptake into dopaminergic neuron, leading to increased cellular damage. In line with this argument, we and others reported previously that in the opposite scenario, overexpression of CAT-2 decreased 6-OHDA-induced dopaminergic neurodegeneration [13,62]. We speculate that *cat-2* deletion might cause different effects in *glit-1* mutants and wild-type animals due to the different concentrations of 6-OHDA used to analyse phenotypes in the respective background: at 0.75 and 10 mM 6-OHDA the detrimental effects of decreased intracellular dopamine levels might be still detectable, whereas at 25 and 50 mM 6-OHDA these effects might not make a difference anymore‥ We cannot explain why mutation of BAS-1 does not lead to similar effects as mutation of CAT-2; however, we note that only CAT-2, but not BAS-1, is specifically involved in dopamine synthesis [23]. Mutation of the vesicle-packing enzyme CAT-1 is in contrast expected to lead to increased cytosolic dopamine. Increased intracellular dopamine levels were suggested to be detrimental for dopaminergic neurons (for review see [63]). We speculate that after exposure of *cat-1* single mutants to high concentrations of 6-OHDA (25 mM and 50 mM), high levels of cytosolic dopamine might buffer the (even more) damaging effects of the drug.

